# A novel method for reliably measuring miniature and spontaneous postsynaptic potentials/currents in whole-cell patch clamp recordings in the central nervous system

**DOI:** 10.1101/2022.03.20.485046

**Authors:** Martynas Dervinis, Guy Major

## Abstract

Measurements of miniature postsynaptic currents (mPSCs) or potentials (mPSPs) in the soma of neurons of the central nervous system (CNS) provide a way of quantifying the synaptic function at the network level and, therefore, are routine in the neurophysiology literature. These miniature responses (or minis) are thought to be elicited by the spontaneous release of a single neurotransmitter vesicle, also called quantum. As such their measurement at the soma can potentially offer a technically straightforward way of estimating ‘quantal sizes’ of central synapses. However, popular methods for detecting minis in whole-cell recordings fall short of being able to reliably distinguish them from background physiological noise. This issue has received very limited attention in the literature and its scope as well as the relative performance of existing algorithms have not been quantified. As a result, solutions for reliably measuring the quantal size in somatic recordings also do not exist. As the first step in proposing and testing a potential solution, we developed and described a novel mPSP/mPSC detection algorithm as part of our quantal analysis software called ‘minis’. We tested its performance in detecting real and simulated minis in whole-cell recordings from pyramidal neurons in rat neocortical slices and compared it to two of the most-used mini detection algorithms. This benchmarking revealed superior detection by our algorithm. The release version of the algorithm also offers great flexibility via graphical and programming interfaces.

## Introduction

Miniature postsynaptic potentials (mPSPs; under voltage recording) or currents (mPSCs; under voltage clamp), often simply called minis, are thought to correspond to the postsynaptic response to a spontaneous release of a single neurotransmitter vesicle (Brown et al., 1979; del Castillo and Katz, 1954; Fatt and Katz, 1952; Isaacson and Walmsley, 1995; Wall and Usowicz, 1998). Somatic measurements of their properties in the CNS provide a window onto the synaptic function at the network level and, therefore, are routine in the literature. Their kinetics (i.e., rise and decay times) are thought to reflect the underlying kinetics of individual ion channels (α-amino-3-hydroxy-5-methyl-4-isoxazolepropionic acid receptors, AMPARs (Sara et al., 2011), or γ-Aminobutyric acid type A receptors, GABA_A_Rs (El Khoueiry et al., 2022); i.e., their probabilities of opening and staying open) (De Koninck and Mody, 1994), as well as the locations of the source synapses across the neuronal dendritic tree (proximal vs. distal) and the passive electric properties of neuronal membranes (Agmon-Snir and Segev, 1993; Major et al., 1994; Rinzel and Rall, 1974). Their incidence rates (often called frequencies, although they do not occur rhythmically) are typically thought to reflect the overall number of active synapses on the dendritic tree (Isaac et al., 1995; Liao et al., 1995; Segal, 2010) but can also be indicative of the rate at which presynaptic vesicles are fusing spontaneously with the neuronal membrane (Glasgow et al., 2019). Finally, the size of the mean amplitude of minis is often thought to reflect the overall strength of synapses across the dendritic tree (Segal, 2010). Assuming that minis are postsynaptic responses to the release of a single neurotransmitter vesicle, their amplitude measurements at the soma can also be used as a straightforward method of estimating the quantal size of synapses onto a recorded neuron. These estimates should, however, be viewed with caution as minis amplitudes at the soma tend to be small and difficult to distinguish from noise in the CNS (Larkum et al., 2009; Major et al., 2013; Nevian et al., 2007; Stuart and Spruston, 1998; Williams and Mitchell, 2008; Williams and Stuart, 2002).

‘Template matching’ and ‘thresholded amplitude detection’ have emerged as the two main approaches to detecting minis (Shi et al., 2010). The first approach relies on the observation that minis often have a fast rise phase and a slower decay phase while noise fluctuation typically do not show such regularities (Liao et al., 1992; Rall, 1967). The somatic current or voltage recording trace is scanned with a sliding window and an average minis template is scaled and fitted to the traces to detect minis-like events of varying amplitudes (Clements and Bekkers, 1997). This approach has been implemented in a commercially available software suite pClamp (Clampfit application). The second approach detects peaks in the recording traces and applies an amplitude threshold combined with shape criteria like the fast rise time and often a limit on the area under the trace (total charge) (Clements and Bekkers, 1997; Shi et al., 2010). This approach has been implemented in a popular commercially available software application MiniAnalysis. ‘Thresholded amplitude detection’ algorithms are thought to suffer from a lack in selectivity (Clements and Bekkers, 1997). As a result, their performance can either inflate false positive detection rates and confuse noise fluctuations with the small amplitude minis if a low amplitude threshold is used or inflate false negative rejection rates and bias the detection towards large amplitude events if a large amplitude detection threshold is used instead. Similarly, change in the signal to noise ratio due to changes in the noise level or the amplitude of minis would artifactually affect estimates of both amplitudes and incidence rates of minis. ‘Template matching’ algorithms were introduced as a remedy to the selectivity problem (Clements and Bekkers, 1997), yet in practice they were found to bias detection towards unambiguous clearcut PSPs that often were also large, and, therefore, they suffer from the opposite problem of the lack in sensitivity (Hwang and Copenhagen, 1999). The lower popularity of ‘template matching’ algorithms compared to their counterpart suggests this problem to be more severe than the one intended to be remedied. Given these considerations and our goal of developing a reliable method for estimating quantal sizes of central synapses based on minis’ amplitude somatic measurements (presented in (Dervinis and Major, 2024)), we developed a novel algorithm, called ‘minis’, to detect miniature (and action potential (AP)-triggered) postsynaptic events. A novel algorithm was needed for several reasons. Firstly, existing algorithms have not been described in sufficient detail in the peer-reviewed literature. Popular algorithms like MiniAnalysis and Clampfit are proprietary. Secondly, because existing algorithms are not open, they are not modifiable. Thus, they cannot be automated and integrated with other software. Lastly, existing algorithms have not yet, to our knowledge, been benchmarked in peer-reviewed articles. To fill these gaps, we developed an algorithm that is transparent, flexible, open-source, and objectively evaluated using standardised criteria.

## Materials and Methods

### Published values for amplitudes and incidence rates of minis

We searched the neuroscience literature between January 1, 2020 – January 31, 2022 for a representative sample of recently reported amplitudes and incidence rate of miniature excitatory postsynaptic currents (mEPSCs) and potentials (mEPSPs) in cortical pyramidal cells, and common data analysis practices in the CNS. We also compared reported mEPSC rates per second to a predicted range of values. We limited our search to publications in English only that reported research results on synaptic function in the CNS. We examined the title and abstract fields of studies in MEDLINE database using the PubMed search engine with the following search keywords: (“spontaneous neurotransmission” OR “spontaneous neurotransmitter release” OR “spontaneous transmitter release” OR “spontaneous postsynaptic” OR “miniature postsynaptic” OR “mEPSP” OR “mEPSC” OR “mPSP” OR “mPSC” OR “mIPSP” OR “mIPSC” OR OR “sEPSP”[Title/Abstract] OR “sEPSC”[Title/Abstract] OR “sPSP”[Title/Abstract] OR “sPSC”[Title/Abstract] OR “sIPSP”[Title/Abstract] OR “sIPSC”[Title/Abstract]) AND (“cortex”[Title/Abstract] OR “cortical”[Title/Abstract] OR “neocortex”[Title/Abstract] OR “neocortical”[Title/Abstract] OR “hippocampus”[Title/Abstract] OR “hippocampal”[Title/Abstract] OR “thalamus”[Title/Abstract] OR “thalamic”[Title/Abstract] OR “brain”[Title/Abstract] OR “cerebral”[Title/Abstract] OR “central nervous system”[Title/Abstract] OR “pyramidal”[Title/Abstract] OR “CNS”[Title/Abstract]). Abbreviations ending in PSP and PP stand for postsynaptic potentials, while PSC and PC stand for postsynaptic currents and should not be confused with end-plate potentials and currents.

### Animals and electrophysiology

All experimental procedures were carried out in accordance with the UK Animals (Scientific Procedures) Act 1986 at Cardiff University under Home Office personal and project licenses. 19 to 27-day old Wistar rats (RRID:RGD_13508588; n = 11) of either sex were anaesthetised with isoflurane and decapitated, and the brain removed under cold (0-2 °C) artificial cerebrospinal fluid (aCSF), composed of (in mM) 125 NaCl, 26 NaHCO_3_, 2.3 KCl, 1.26 KH_2_PO_4_, 1 MgSO_4_, 10 glucose, and 2 CaCl_2_ bubbled with 95% O_2_ 5%CO_2_. Coronal somatosensory cortical slices 350 µm thick were cut and held in an interface chamber with their surfaces kept moist for 1 to 3 hours by aCSF at room temperature (21-23°C) under a 95% O_2_ 5% CO_2_ atmosphere. Individual slices were then placed in the recording chamber and held at 35-37°C with aCSF flowing over both surfaces. For recording purposes at 35-37°C the aCSF solution contained (in mM) 125 NaCl, 24 NaHCO_3_, 2.3 KCl, 1.26 KH_2_PO_4_, 1 MgSO_4_, 10 glucose, and 2 CaCl_2_.

Whole-cell recordings (n = 14) were obtained under visual control using infrared scanning gradient contrast imaging with the imaging laser beam and a Dodt tube. Recordings were made with 4.5-9 MΩ borosilicate glass pipettes (Harvard Apparatus, 1.5 mm outside diameter and 0.86 mm, internal diameter, omega-dot filament; Sutter P2000 puller), filled with (mM): 140 KGluconate, 10 HEPES, 2 MgCl_2_, 3 ATP-Na_2_, 0.3 GTP-Na, 0.1 Magnesium Green, 0.1 Alexa 594; 7.35 pH, 273 mOsm/L osmolarity. Signals were amplified, low-pass filtered (5 KHz) and digitised (Axopatch-200B, custom amplifier/filters, Digidata 1440, Clampex software; Molecular Devices). Pipettes and electrodes were positioned with Sutter MP265 and 285 manipulators with a diagonal (axial) mode, using parallelogram trajectories.

The electrophysiological recordings consisted of two phases. During the first phase we recorded mEPSPs against the background of physiological noise (‘noise with minis’ condition) by applying aCSF with AP and inhibitory postsynaptic potential blockers added: 0.5 or 1 μM tetrodotoxin (TTX; selective, potent Na^+^ channel blocker), and 12.5 μM gabazine (selective, potent GABA_A_ receptor antagonist to block (fast components) of inhibitory synaptic inputs – IPSCs and resulting IPSPs). AP blockade by TTX was explicitly tested for with long current pulses pushing membrane potential (V_m_) well above normal AP threshold levels, to around –20 mV. The average duration of this phase of the recordings was 21.8 (± s.d. 0.9) minutes. During a second phase, we recorded noise with mEPSPs blocked (‘noise-alone’ condition). This recording was carried out after further adding 40 μM 2,3-dioxo-6-nitro-7-sulfamoyl-benzo[f]quinoxaline (NBQX) (selective, potent α-amino-3-hydroxy-5-methyl-4-isoxazolepropionic acid (AMPA) receptor antagonist) and 50 μM CPP (competitive, non-use dependent selective N-methyl-D-aspartate (NMDA) receptor antagonist) to the solution used in the ‘noise with minis’ condition, to block all excitatory glutamatergic synaptic inputs (and thus EPSPs), verified by inspecting and analysing the traces across a number of recordings with the same flow rates. The average duration of this phase of the recordings was 44.8 (± s.d. 2.9) minutes.

### Experimental design

The goal of this benchmarking study was to compare mEPSP detection performance by three different detection algorithms, mainly in terms of standard signal detection theory measures. Two of them were the most popular software products used in the field of synaptic function research in the CNS, while the third one was our novel algorithm called ‘minis’.

Current clamp whole-cell patch recordings obtained during the initial recording phase (‘noise + minis’ condition) underwent mEPSP detection analysis using ‘minis’, MiniAnalysis (Bluecell; RRID:SCR_002184), and Clampfit (part of the pClamp software suite, Molecular Devices; RRID:SCR_011323). Detected mEPSPs were analysed in terms of their amplitude, 10-90% rise and decay times, and incidence rate.

The current clamp recordings obtained during the second recording phase (‘noise-alone’ condition) served as the background data in the mEPSP (‘simulated minis’) computer simulations (n=14 cells). Computationally simulated mEPSPs (smEPSPs) were added onto segments of these V_m_ noise recordings (1 of them 200 seconds, 1 more of them 40 seconds, while the remaining 12 were 100 seconds long) and the resultant hybrid waveforms (noise with simulated minis) were then subjected to mEPSP detection analyses using ‘minis’, MiniAnalysis, and Clampfit software. Detection performance was quantified using signal detection theory measures and compared between the three algorithms.

### Simulations

#### Simulating distributions of miniature excitatory postsynaptic potentials with a range of amplitudes and rise times

All simulations were carried out within the ‘minis’ software environment which is based on Matlab (Mathworks; RRID:SCR_001622). Simulation of minis-like events was based on an analytical solution to the passive cable equation for a transient current (Rall, 1977) using 100 lumped terms with a double exponential synaptic current with τ_1_= 0.1 ms and τ_2_ = 2 ms (Major et al., 1993).

For a given membrane time constant, an initial cable segment was constructed with an electrotonic length *l*/λ = 0.6, where *l* is the physical length of a dendrite (μm) and λ is the length (μm) constant (Rall, 1977). Charge (Q) was injected at a location (x μm) along this cable and the resulting simulated postsynaptic V_m_ was measured at one end of this cable segment. The precise location of the injected charge was varied systematically, depending on the desired shape of the simulated event. Increasing the distance of the injected charge along the cable away from the measurement site gradually increased the rise time of the simulated event. To further increase the rise time, electrotonic length was gradually increased until the required rise time was obtained. Increasing the amount of injected charge increased the amplitude of the simulated event. Therefore, one was able to construct a simulated postsynaptic potential of any rise time or amplitude, for the particular membrane time constant used. A pool of simulated events could be created having any kind of distribution of amplitudes and rise times, for a given decay time constant, with individual events’ onsets timed/placed pseudo-randomly along the time axis of the V_m_ recording.

#### Simulations used to construct full receiver operating characteristic (ROC)-like curves, using relatively fixed but realistic (‘moderately-sized’) mini amplitudes

All simulated mEPSPs were drawn from a two-dimensional normal distribution (independent amplitude and rise time dimensions) with the following amplitude parameters: mean μ_1_ = 0.3 mV and standard deviation σ_1_ = 0.05 mV (i.e., amplitudes relatively constant, with a relatively small amount of variation). The following (independent) 10-90% rise time parameters were used: μ_2_ = 0.05 ms and σ_2_ = 2.5 ms (with a minimum rise time of 0.05 ms; i.e., only upper half of normal distribution used). Simulated events were drawn pseudo-randomly from a distribution at one of the following incidence rates (minis/second): 640, 320, 160, 80, 40, 20, 10, 5, 2.5, 2.5, 2.5, 2.5, 2.5, and 2.5. Corresponding noise amplitude scale factors for these 14 incidence rates conditions were as follows: 1, 1, 1, 1, 1, 1, 1, 1, 1, 1.2, 1.4, 1.8, 2.6, and 4.2. The simulated events were then positioned at pseudo-randomly determined times along the V_m_ noise recording epochs obtained during the second recording phase (‘noise-alone’ condition). Two of the noise-alone epochs were 200 and 40 seconds long, while the remaining twelve were 100 seconds long. Four different simulation traces per recording, each of them having different pseudo-random event timings, were generated for every incidence rate/noise scale condition, resulting in a total number of 14×4×14 = 784 traces.

#### Simulations of miniature excitatory postsynaptic potentials using a range of biologically realistic amplitudes at realistic incidence rates

The simulation parameters used to construct full ROC-like curves cover only a part of the likely biological range. In reality, in cortex, minis vary far more in their amplitudes, with large numbers of smaller minis, but a minority of bigger minis too – some much bigger. Therefore, we set out to compare all three algorithms using a range of plausible simulated mini amplitudes, within the restricted range of what we deemed to be realistic incidence rates based on our estimation (from recent peer-reviewed publications) in the first Results subsection.

All simulated events were drawn from a two-dimensional distribution with the following parameters: amplitudes were chosen from ten discrete values (0.05, 0.1, 0.15, 0.2, 0.25, 0.3, 0.35, 0.4, 0.45, 0.5 mV) with equal probabilities, rise times were independently chosen from the upper half of a normal distribution with mean μ_2_ = 0.05 ms, and standard deviation σ_2_ = 2.5 ms (i.e., no rise time shorter than 0.05 ms). Three ‘realistic’ minis’ incidence rates were used: either 61, 38, or 27 minis/s. The simulated events were positioned at pseudo-random times along the same noise recording epochs as used in the previous subsection. Four different simulation traces per recording, each of them with different random event timings, were generated for every incidence rate condition, resulting in a total number of 3×4×14 = 168 traces.

### Detection of spontaneous and miniature postsynaptic potentials

#### ‘minis’ detection algorithm

We developed a new algorithm to detect postsynaptic potentials and currents, that we have incorporated into a data analysis software package called ‘minis’, coded in Matlab (Mathworks). It is distributed as an application programming interface (API) in the form of a packaged Matlab app or a Python (Python Software Foundation; RRID:SCR_008394) package or as a compiled standalone desktop application with a graphical user interface. In brief, the detection algorithm takes a filtered V_m_ or clamp current trace in the Axon Binary File (ABF) format and detects peaks and rising phases and estimates their amplitudes and rise and decay times after applying certain processing steps.

These steps are outlined below (with parameter values tailored for detecting smEPSPs in cortical pyramidal neurons). A schematic illustration of the key steps is shown in Figure 1:

A. Steps to find minis:
  1. The data can be band-stop filtered to remove mains noise (e.g., small integer multiples of 50 Hz frequency components: optional). For this, a Butterworth filter (Matlab’s ‘butter’ function) is used with a stopband attenuation of 10 dB and a passband ‘ripple’ of 0.05 dB. The stopband size is 1 Hz, and the passband is the entire remaining frequency range except for the ±3 Hz window either side of the stopband frequency. Filtering effects are illustrated in Figure 2.
  2. The recording trace can be smoothed using a Gaussian window (optional). We use a 1.5-millisecond window (with standard deviation of the Gaussian of 0.53 ms). This step removes high frequency noise. Smoothing effects are illustrated in Figure 2.
  3. Peaks (local maxima) are identified in the filtered, smoothed recording trace. Each peak is then examined consecutively, starting with the initial peak.
  4. Going through all the peaks one by one, any peak within the “Peak Integration Period” (‘duration’) following the current (‘working’) peak of interest is discarded if it is *smaller* (lower) than the peak of interest. For the analysis in this paper, we use a Peak Integration Period of 2.5 ms; this can be matched to the different ‘sharpnesses’ or ‘breadths’ of peaks in different data.
  5. If there is a larger peak within the peak integration period, the ‘working’ peak of interest is moved to this new peak, and the old peak and all intervening peaks between it and the new ‘working’ peak of interest are discarded.
  6. The baseline of the peak is positioned asymmetrically around the lowest value (local minimum) before the peak. Because of the faster upstroke of minis, compared with their decays, 80% of the baseline period falls before the lowest voltage (local minimum) and the remaining 20% after it (see Figure 1, inset). The interval from the baseline to the peak is limited by the *Maximum Time-to-Peak* parameter (10 ms used here). The last 20% of the baseline cannot start earlier than this.
  7. The length in time of the baseline is controlled by the *baseline duration* parameter. We used 2-ms. (N.B. a baseline, or indeed an entire mini, can occur during the summated decay ‘tails’ of previous minis, particularly when the minis incidence rate is high relative to their decay time, as often seen with mPSPs under voltage recording at body temperature – but also under voltage clamp).
  8. If the baseline overlaps with or falls before the previous peak, its start is delayed till immediately after the previous peak (to stop the baseline being pulled down by the previous rising phase), but the end time is kept the same.
  9. To reject any large artefactual ‘glitches’, or small events that are putative noise, if the amplitude of the peak is outside of the range of acceptable amplitudes, between a detection threshold and a maximum, the peak is discarded. We allowed a range of amplitudes of 0.1-10 mV in detection conditions used to construct full ‘virtual’ receiver operating characteristic (ROC) curves. In other conditions we applied a range of 0.05-10 mV. To detect real minis, we used a range of 0.01-10 mV. (N.B. 0.02 or 0.03 mV detection thresholds can also be used for real whole-cell patch recording data from neocortical pyramidal neurons, under our experimental conditions, guided by the range of amplitudes of dendritically detected, somatically recorded mEPSPs in (Nevian et al., 2007) from layer 5 pyramidal neurons from rat somatosensory cortex; in general, detection thresholds will need to be adjusted to the cell type and experimental conditions).
B. Steps against multiple small minis being erroneously lumped together (merged):
  10. Rising phase asymmetries: if the first half of the rise time (10-50% or 20-50%, depending on the rise time duration of choice) is longer than the second half (50-90% or 50-80%) by a factor of 5 or more (an asymmetry incompatible with typical single, non-summating EPSP or EPSC rising phases), the end of the baseline is extended slightly into the ‘take-off’ of the rising phase. (This correction can be repeated one more time).
  11. The amplitude range test is applied again (Step 9).
  12. If the second half of the rise time is longer than the first half by a factor of 5 or more, the peak is shifted to a previously discarded smaller peak, if exists. (Two minis could be summating on their rising phases, which is treated as a single mini, or the second mini could occur just after the peak of the first, which is treated as two minis, if two peaks are distinguishable).
  13. The amplitude range test is applied again (Step 9).
  14. If the 10% or 20% rise time mark (depending on whether 10-90% or 20-80% rise time is measured) is later than the end of the baseline by more than a half of the baseline duration (1 ms in our case), a new baseline is established closer to the peak (by moving the baseline to the next local minimum).
  15. Finally, the amplitude range test is applied again (Step 9).
  16. A mini (mPSP/mPSC) is established.

**Figure 1:**
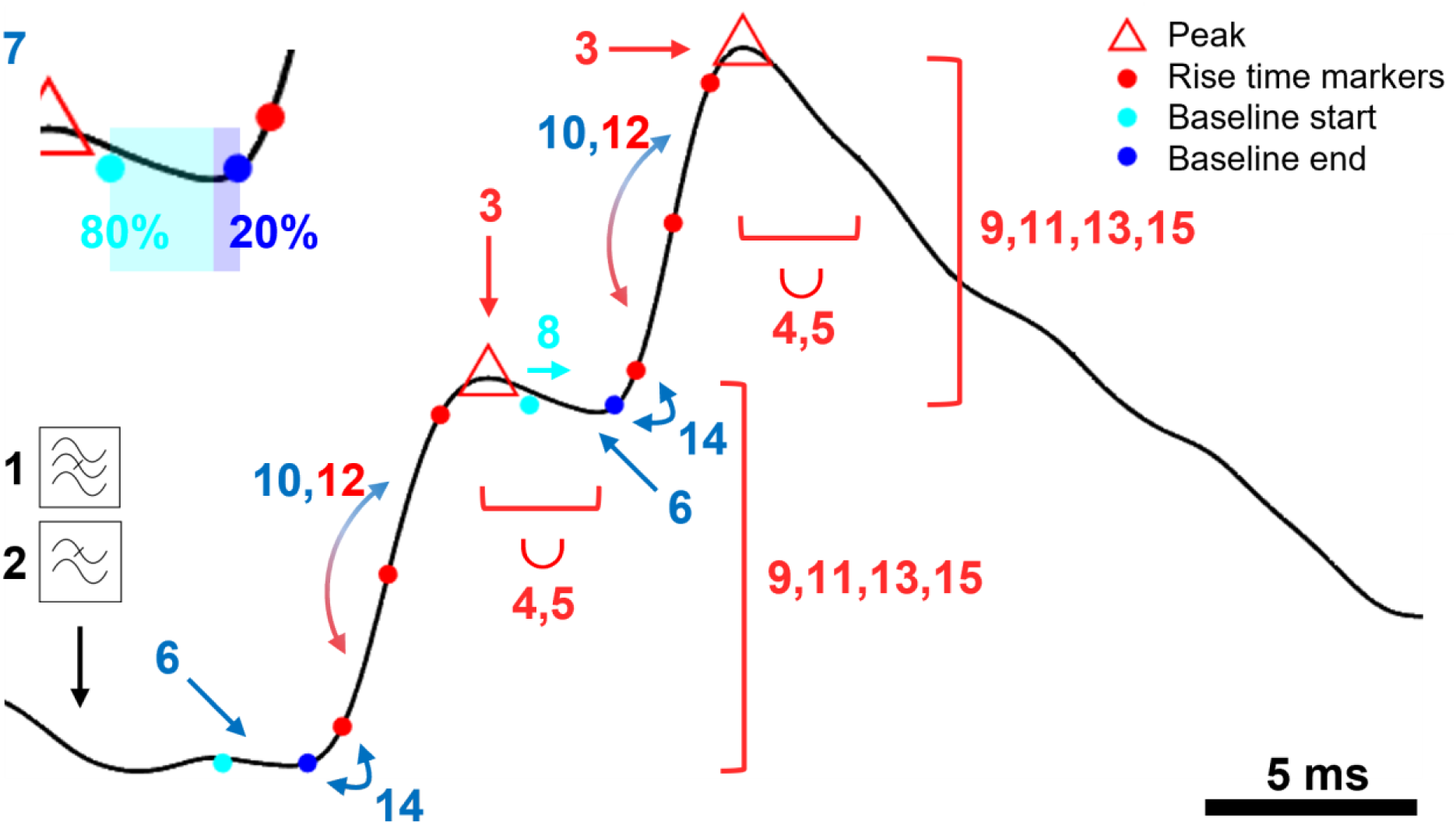
A Schematic representation of key steps in ‘minis’ detection algorithm. An illustration of a step-by-step application of the mPSP/C detection algorithm using simulated data. Numbered steps dealing with mini peaks/amplitudes are shown in red, steps used to determine the baseline are shown in blue. Briefly, the steps are: 1. Bandstop filter (against mains). 2. Lowpass filter to remove high frequency noise. 3. Find peaks. 4. Discard smaller peaks with the peak integration period represented by the length of the bracket and the union symbol (∪). 5. Move to a higher peak (if exists) within the peak integration period discarding the earlier peak. 6. Position the baseline at a local minimum located within the time-to-peak period prior to the peak. 7. Position the baseline asymmetrically (80:20) around the minimum (colour change at the inset). 8. Correct the start of the baseline if it overlaps with the previous peak. 9. Discard the peak if it falls outside the acceptable amplitude range (repeated three more times). 10. Compare rise time halves to correct the baseline (can be repeated once more). 11. Repeat Step 9. 12. Compare rise time halves to correct the peak. 13. Repeat Step 9. 14. Correct the baseline if it significantly deviates from the 10%/20% rise time mark. 15. Repeat Step 9.

**Figure 2:**
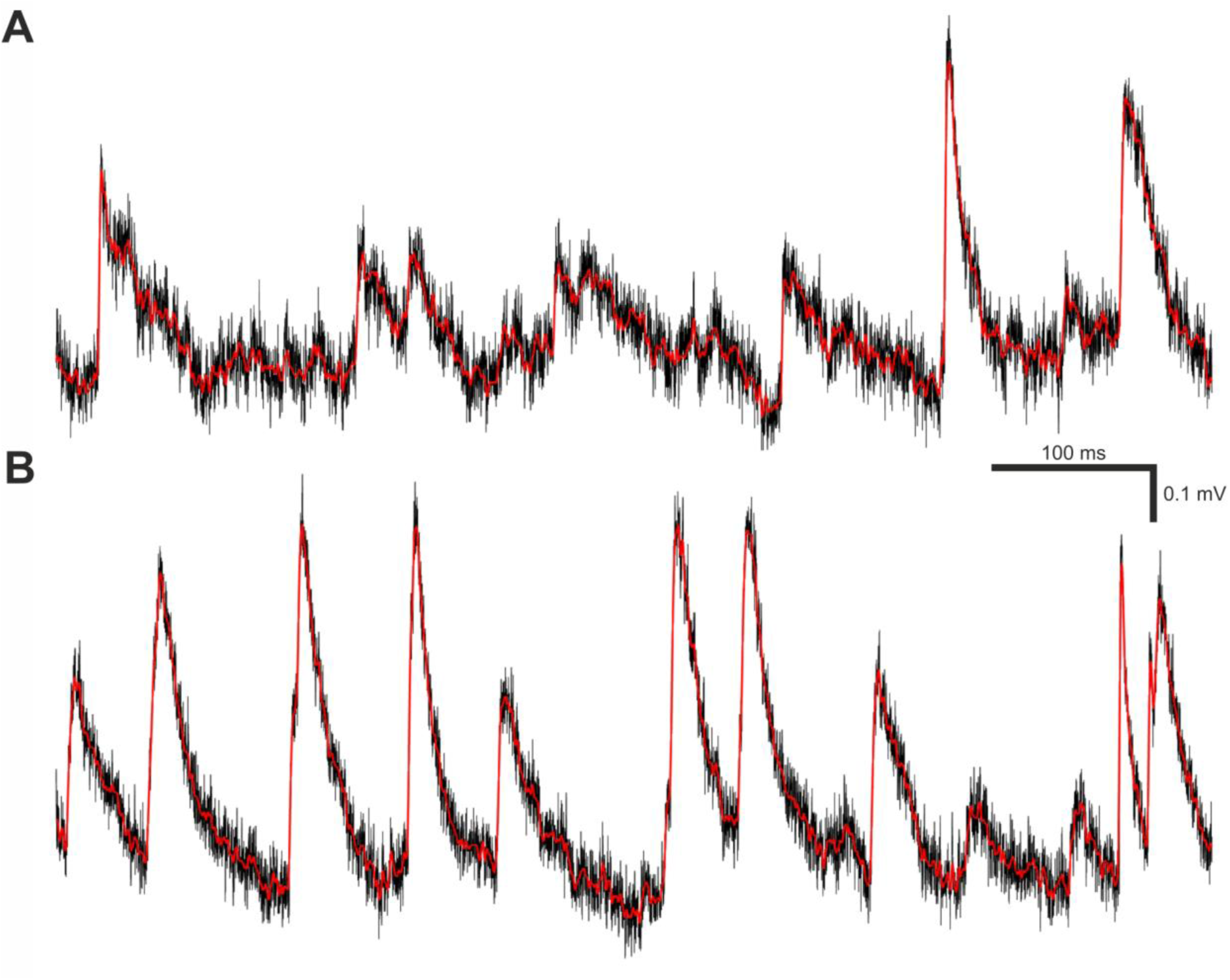
Filtering and smoothing effects. (A) A brief segment of V_m_ trace recording containing real minis and noise. The black trace is raw (unfiltered and unsmoothed), whereas the red trace is the same data after filtering and smoothing as described in Materials and Methods subsection called *‘minis’ detection algorithm*. (B) Same as (A) but containing varying amplitude smEPSPs with a 20 minis/s incidence rate instead of real minis.

#### Detection using MiniAnalysis software

MiniAnalysis software version 6.0.7 was used to detect mEPSPs for a benchmarking purpose. We used the following detection parameters: As with the ‘minis’ program, an amplitude threshold of 0.1 mV (when constructing the full ROC-like curve) or 0.05 mV (all other simulated conditions), in both cases somewhat matched to the simulated data, or 0.01 mV when detecting real minis, a “Period to search for a Local maximum” of 12.5 ms, “Time before a peak for baseline” of 10 ms, a “Period to search a decay time” of 30 ms, a “Fraction of peak to find a decay time” of 0.37 (one effective time constant for exponential decay), a “Period to average a baseline” of 2 ms, an “Area threshold” of 0 mV×ms (i.e., not used), a “Number of points to average for peak” of 1, and using the “Detect complex peak” option (to avoid missing close-together events). The chosen detection parameters closely resembled similarly named parameters in the ‘minis’ detection algorithm with minor deviations to improve performance. Arrived by trial and error, these parameters were also found to give the most optimal detection performance of the MiniAnalysis software in comparison to other tested parameter sets. Event detection was carried out on the recording data that was filtered and smoothed in the same way as described in steps 1 and 2 of the previous sub-section (Figure 2).

#### Detection using Clampfit

Clampfit software, part of the pClamp 11.0.3.03 software suite, was used to detect real and simulated mEPSPs and mEPSCs for a benchmarking purpose. Recordings were band-stop filtered as described in step 1 of the ‘minis’ detection algorithm. Data was not smoothed. A template search algorithm was used with 9 templates. The mEPSP template set was constructed based on the simulated events. Events having 0.5 ms, 1.5 ms, 3.5 ms, and 6.5 ms 10-90% rise times were selected to produce 4 distinct mEPSP templates (with the correct decay time constants). An additional 5 single-peaked templates were constructed in a way that made them potentially a mix of multiple simulated waveforms. They were constructed to have extra-fast, fast, intermediate, slow, and extra-slow rise times corresponding to 2 ms, 2.75 ms, 6 ms, 7 ms, and 8 ms 0-100% rise times and 8 ms, 9.25 ms, 17 ms, 23 ms, and 37 ms duration full decays, respectively. Combining single and mixed waveforms gave better detection performance than having waveforms based only on any one of these types alone. Template match threshold was set to the default of 4 noise standard deviations with the rest of the detection parameters being set to their default values as well. The same set of templates was used to analyse all simulated current clamp data, as well as real data. Using the same templates with real data was justified, because they resulted in a superior detection performance compared to templates created directly from the real data.

#### Detection algorithm response classification

At the outset we identified prominent noise events in the V_m_ noise recordings (the 2^nd^ recording phase: ‘noise-alone’ condition) that could potentially be confused for true minis (mEPSPs) by any of the algorithms (noise traces in Figures 3A and B). All noise recordings were filtered and smoothed as in Steps 1 and 2 of the ‘minis’ detection algorithm (Figure 2). The data was further smoothed by a rectangular ‘box-car’ moving average window of 20 ms. Then all peaks larger than 0.01 mV and having a half-width of at least 0.5 ms were identified. These peaks were classed as (notional) noise ‘events’ (for the purpose of generating ‘sensible’ true negative rates, TNRs, which do not ‘swamp’ false positive rates across the entire virtual ROC; see below). Subsequently we simulated mEPSPs and added them at pseudo-random times to a zero-trace having the same length as the noise recording of interest (Simulation traces in Figures 3A and B). The peaks of these events were classed as signal events. The simulated trace was then added to the noise recording (“Simulation + noise” traces in Figures 3A and B). Locations of signal and designated noise events formed the ‘ground truth’ information.

**Figure 3:**
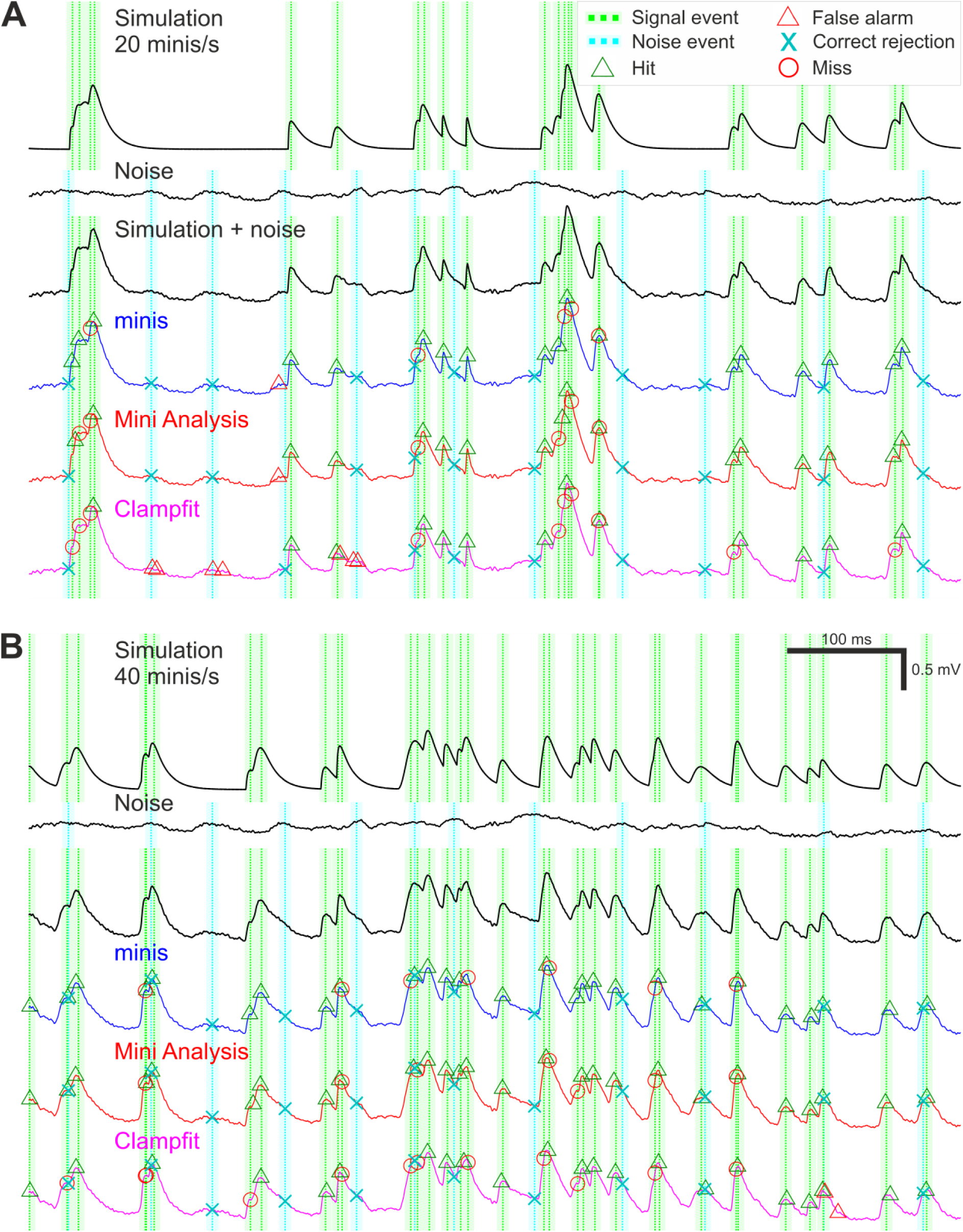
Simulation and detection performance. (A) An example of simulated membrane potential recording with average simulated mEPSP incidence rate of 20 minis/s. The top voltage trace shows a brief segment with randomly timed smEPSPs with random, normally-distributed amplitudes (μ_1_= 0.3 mV, σ_1_ = 0.05 mV) and 10-90% rise times (drawn from the upper half of the distribution with μ_2_ = 0.05 ms, σ_2_ = 2.5 ms); see Methods. Dotted green vertical lines mark signal events corresponding to smEPSP peaks. The paler shaded colour demarcates a 10-ms (± 5 ms) window for accepting the detection of a signal event. The second traces from the top are corresponding recordings of V_m_ noise fluctuations (the 2^nd^ recording phase: ‘noise-alone’ condition, in TTX and transmitter blockers), after initial filtering and smoothing (steps 1 and 2 of the ‘minis’ detection algorithm; Figure 1). Dotted vertical cyan lines indicate noise events; the shaded colour demarcates a window of exclusion for correctly rejecting these events (see Methods). The V_m_ third from the top trace is a hybrid one with smEPSPs added to the noise recording and with signal events and noise ‘events’ indicated. The coloured V_m_ traces below show detection performance for the three different algorithms. (B) An example of simulated V_m_ recording with mEPSP incidence rate of 40 minis/s.

Following the detection process, all detected events were associated with one of the signal and noise events depending on their proximity to these predefined ‘ground truth’ events. A detected event was classed as a true positive (a ‘hit’) if it was the closest detection event within 5ms of a signal (simulated mEPSP/mini) event. If no detection event occurred within the 10 ms symmetrical window surrounding a signal (simulated mini) event (–5 to 5 ms), the signal event was classed as a false negative (a ‘miss’). If no detection event (except correct detections) occurred within the same duration window surrounding a noise event, the noise event was classed as a true negative (‘correct rejection’). Although in theory there could potentially be a much higher number of possible time points which could be labelled as correct rejections, or true negatives, in this situation, because of their inherently somewhat arbitrary nature, we wanted to keep the maximum number comparable to the maximum number of true positives (simulated minis), so the ROC plot had a reasonable chance of being relatively ‘square’ (i.e. having a useful dynamic range). The remaining detection events were classed as false positives (‘false alarms’). Examples of classified detection events are shown in Figure 3 separately for all three detection algorithms.

#### Detection response time to the nearest neighbour

The time to the peak of the nearest smEPSP was available for every classified response. Detection performance was then evaluated as a function of the time to the nearest neighbour.

### Membrane potential ‘phase’ (trend) classification

We attempted to address common detection errors, in particular missed minis on the decaying phase of a previous mini (causing second peak to drop relative to its baseline), or on the rising phase of another mini (so the rises of both minis get lumped together). Algorithm detection performance was assessed not only for full recordings but also separately for periods when the ‘underlying’ simulated V_m_ (before adding the noise) was rising, decaying, or remained roughly stable (Figure 4). The ‘decaying phases’ (downward trend) of the purely simulated V_m_ were set to be a) the period from 0.0625 ms after the peak until the purely simulated V_m_ trace dropped to the ratio of 0.3/e mV (0.11 mV) (amplitudes of simulated minis were ∼0.3 mV in the full virtual-ROC curve conditions, so this would be about one membrane time constant after an ‘average’ simulated mini) or b) until the V_m_ started rising again to reach a new peak that was above the initial peak (at the start of this decay phase). If the new peak was lower than the initial one, the entire new peak was classified as part of the ongoing decay phase. Peaks occurring on the decay phase of the V_m_ were assigned the V_m_ decay rate of change that was present 0.25 ms (5 sample points) prior to the appearance of the V_m_ inflection points associated with the start of the rise of a new mini (indicated by the blue arrow in Figure 4A). The rising ‘phase’ was treated essentially somewhat as a decay phase in reverse. The only difference was that the ratio of 0.3/e mV was replaced by the ratio of 0.3/10 mV (0.03 mV). Peaks occurring on a V_m_ rise phase (upward trend) were assigned a V_m_ rate of rise value that was present 0.125 ms *after* the appearance of an upward V_m_ inflection point following the initial decay of the previous peak (indicated by green arrows in Figure 4A). Periods that were already classified as decay phases could not be reclassified as rising phases. Periods of relatively slowly-changing simulated V_m_, outside of the rise and decay phases were classed as “stable” (or a flattish phase). The classification of purely simulated membrane potentials into three different trend phases was used to assign phases to the combined noise recordings with simulated minis (Figure 4B). The detection performance during rise and decay trend phases was evaluated as a function of the V_m_ rate of change.

**Figure 4:**
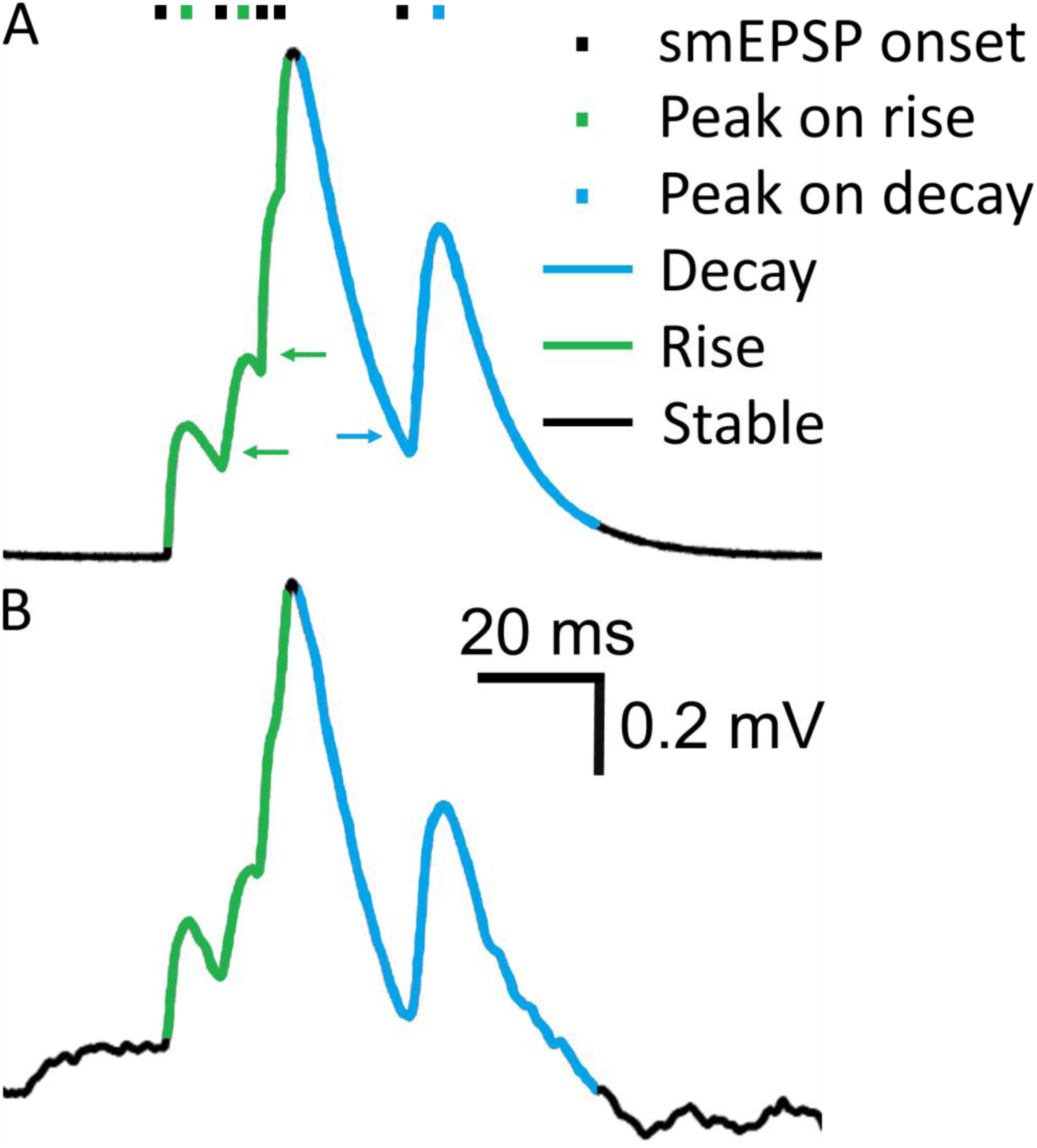
Membrane potential rising, steady-ish and falling ‘trend phases’. (A) A short segment of a V_m_ trace showing five smEPSPs. Different V_m_ ‘trend phases’, based on the overall upward or downward trend in *simulated* voltage, are marked in colour. Green arrows indicate V_m_ points where the V_m_ rate of increase is assigned to the preceding mEPSP peak. The blue arrow indicates a V_m_ point where the V_m_ rate of decay is assigned to the following mEPSP peak. (B) Simulated V_m_ trace in (A) added to a noise recording. V_m_ ‘trend phases’ were determined in (A) and are used to classify detected events in (B).

## Data Analyses

Performance of the three algorithms was compared using measures from signal detection theory. All detection responses were classified as described in the *Detection algorithm response classification* subsection. True positive (‘hit’) rate (TPR) was calculated as follows:

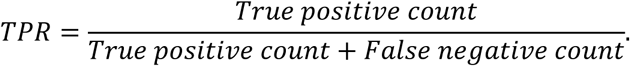

Throughout this text we use TPR, detection probability, and sensitivity interchangeably. False positive (‘false alarm’) rate (FPR) was calculated as follows:

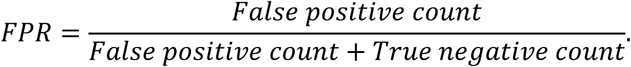

The true negative count we have used is perforce notional, as explained above, but still useful for standardised, non-subjective comparisons and benchmarking. Throughout we use FPR and ‘false alarm rate’ interchangeably.

Calculation of TPR and (notional) FPR allowed us to produce ROC-like curves and to calculate area under the curve (AUC; area under the ROC curve, for each complete ROC curve, with 0.5 indicating chance performance (the area of the triangle under the diagonal from (0,0) to (1,1) and 1 indicating perfect performance, i.e., the area of a right angle starting at (0,0), vertical to (0,1) then going horizontal to (1,1)) as detection performance indicators. These are the most commonly used performance indicators for classifiers, in addition to sensitivity (TPR), and d’ (discriminability or sensitivity index) which we often use in parallel throughout this text. The d’ statistic was calculated as follows:

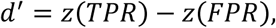

where z is the z-score defined as the inverse of the normal cumulative distribution function, i.e. z(arg) is the number of sd’s from the mean giving the area equal to its argument arg under the standard unit normal distribution (mean 0, sd 1), starting from –∞. TPR and FPR values are essentially treated as cumulative probabilities as explained in (Macmillan and Creelman, 2005). So, d’ tells us how many standard deviations ‘apart’ the TPR and FPR are, assuming they are cumulative probabilities (areas under curve) of a standard normal distribution. N.B. If FPR is 0, z(FPR) is –∞ so d’ is undefined, but arbitrarily large and positive (as occurs in some of our figures below). These plots are truncated for the detection method in question, but the performance is still superior to any plots with finite values.

The above detection performance indicators were calculated for full recordings, as well as separately for rising and decaying periods of V_m_.

In addition, we also looked at cumulative (left-to-right over the x-axis) TPR and cumulative FPR. The cumulative TPR was defined as follows, for each detection algorithm and simulated waveform:

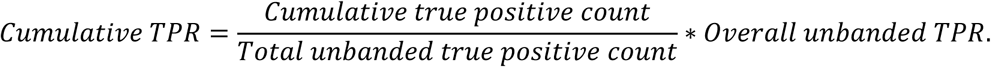

Likewise, the cumulative FPR was defined as:

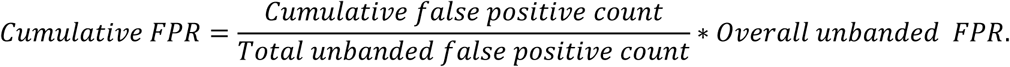

The latter two measures were not only useful for measuring cumulative hits and false alarms over a range of time intervals and V_m_ rates of change, but also provided overall estimates of TPR and FPR (the right side of the x-axis).

## Statistical analyses

All inferential statistics were based on the assumption that distributions of all measures were normal. Only repeated samples or single-sample (often when comparing with baseline true positive values in tables) t-tests were used across the Results section. All mean distribution values are stated/depicted with 95% confidence limits of the mean (if exist).

## Data accessibility

All data analysed in this study is publicly available (Dervinis, 2024a). The available data include ‘noise with minis’ and ‘noise-alone’ whole-cell patch membrane potential recordings that were used in minis detection procedures and computer simulations. The hybrid V_m_ traces containing smEPSPs added to noise recordings are also available. All electrophysiological recordings were stored in Axon Binary File (ABF) format.

mEPSP templates used for detecting real and simulated events via Clampfit are also in the same dataset in the form of Axon Template Files (ATFs).

## Code accessibility

All analyses were carried out in Matlab and the analysis code is publicly available on GitHub (Dervinis, 2024b). The code is complete with instructions on how to reproduce all figures reported in this study.

## Software accessibility

The present study reported the use of a novel mEPSP/mEPSC detection algorithm that is part of ‘minis’ software available on GitHub (Dervinis, 2024c).

## Results

### Research literature measuring minis in the central nervous system

The full results of the literature survey are reported in Supplementary Results. Here we only briefly report key findings. First, we established that the two most popular software applications used to detect minis are MiniAnalysis followed by Clampfit with 48% and 26% of surveyed studies reporting to use these algorithms, respectively. Moreover, these studies reported a mean minis’ incidence rate of 3.6 ± s.d. 0.34 per second (range of 1.0 to 8.0 per second). Meanwhile, we suspect that true minis incidence rates range somewhere between ca. 10 minis/s and 100 minis/s based on research on synapse counts and release probabilities in the CNS (an order of magnitude higher than reported direct measurements using voltage clamp).

### Algorithm performance comparison for detecting moderately-sized (∼0.3 mV) simulated miniature excitatory postsynaptic potentials

#### Detection under a wide range of incidence rate conditions: overall performance

In the Results subsection ‘Research literature measuring minis in the central nervous system’, we reported that detection of spontaneous postsynaptic events is typically carried out using an amplitude threshold. This approach implies that the range of amplitudes of spontaneous postsynaptic events is somewhat known and that, on the smaller end, the amplitudes are moderately-sized and sufficiently larger than pure noise fluctuations. Leaving aside the question of the empirical validity of this (somewhat dubious?) assumption, we evaluated automated detection of moderately-sized (0.3 ± s.d. 0.05 mV) smEPSPs added to whole-cell patch clamp noise recordings (the 2^nd^ recording phase: ‘noise-alone’ condition).

We compared our novel ‘minis’ algorithm with MiniAnalysis and Clampfit. We created 14 simulation incidence rate conditions with different effective signal-to-noise ratios that loosely mimicked the effects of varying the signal detection threshold. On the one hand, increasing the incidence rate of smEPSPs tended to decrease TPR as simulated events started to merge. On the other hand, increasing the noise amplitude (scaling up the noise that was added to the simulated minis) tended primarily to increase the FPR.

We plotted the detection performance in terms of TPRs against FPRs, as ‘virtual’ (notional) Receiver Operator Characteristic (v-ROC) curves, for all 14 conditions and all three algorithms in Figure 5A. We found that in the first six conditions (high minis incidence rates), the v-ROC curve for ‘minis’ had minimal overlap with any other curve and was positioned furthest away from the diagonal (starting from left to right; 640, 320, 160, 80, 40, and 20 minis/s with x1 noise scaling), indicative of a superior performance in this (relatively high) incidence rate range (the highest incidence rates were used to create the full v-ROC; higher incidence rates increase overlaps, i.e. temporal summation, between minis, which are particularly challenging to detect). The same observation was supported by another measure, the sensitivity index d’, shown in Figure 5B. Detection performance in the remaining 8 conditions (10, 5, 2.5, 2.5, 2.5, 2.5, 2.5, and 2.5 minis/s with noise scaling factors of x 1, 1, 1, 1.2, 1.4, 1.8, 2.6, and 4.2, respectively) was very similar for both ‘minis’ and MiniAnalysis: neither of the two showed a clear superiority on both performance measures simultaneously: v-ROC and d’ curves overlapped. In contrast, Clampfit consistently performed worst across all 14 conditions, particularly with high amplitude noise. Clampfit becomes progressively ‘confused’ as noise increases, placing templates on noise events, but ignoring ‘swamped’ smEPSPs.

**Figure 5:**
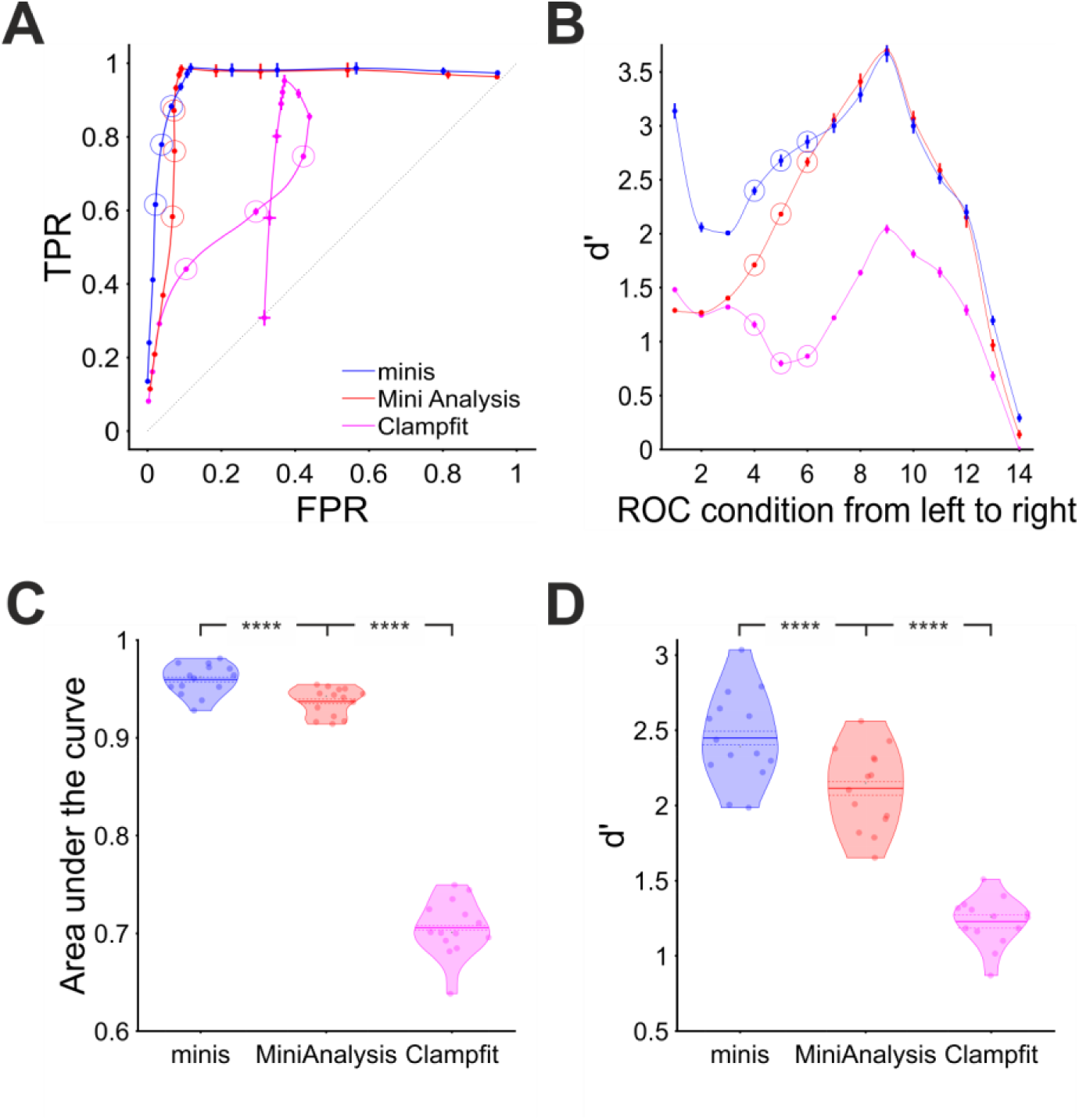
Overall performance for detecting ‘moderately’-sized smEPSPs (∼0.3 mV). (A) ‘virtual’ ROC (Receiver Operator Characteristic)-like curve showing performance for detecting ‘moderately’-sized smEPSPs in terms of True Positive Rate TPR (sensitivity) and False Positive Rate FPR (1 – specificity) for all three algorithms, averaged across all 14 cells. In each case, dataset varies from point to point, as opposed to detection threshold itself (which is varied systematically in a traditional ROC, using one dataset). Ideal performance would be a right-angle plot going vertically up TPR-axis at FPR=0, then horizontally along the top of the plot at TPR=1; ‘minis’ is close to ideal, and superior to MiniAnalysis, which is not too far from ideal (despite its ‘missed minis’ problems). Clampfit is far from ideal. Different points represent 14 different incidence rate/signal-to-noise ratio conditions, averaged across all 14 cells. Moving (generally) from bottom to top/left to right, simulated minis rates and signal-to-noise ratio are decreasing. Vertical and horizontal bars indicate 95% confidence interval around each mean. The dotted diagonal line indicates chance performance. The hollow circles mark three realistic smEPSP incidence rates (80, 40, and 20 minis/s). (B) Sensitivity (discriminability) index (d’) in the same 14 conditions as in (A) for all three algorithms, averaged across all 14 cells. (C) ‘Violin’ plots for Area under the virtual-ROC curve in (A) for all three algorithms. Individual data points represent the 14 individual recordings (cells), horizontally ‘swarmed’ to avoid overlaps. AUC (Area Under Curve) is averaged across all 14 incidence rate/signal-to-noise conditions for each cell for each detection algorithm. The mean is marked by a solid line over the ‘violin’ centre. The dashed line indicates the 95% confidence limits of the mean, and coloured shapes indicate the approximate (smoothed) probability densities (wider = higher probability of those values). **** indicates (highly) significantly different at p ≤ 10^-5^ level, paired t-test. (D) Similar plot for sensitivity index d’ averaged across all 14 incidence rate/signal-to-noise conditions for each cell and each detection algorithm.

When overall performance was assessed averaging across individual incidence rate and signal-to-noise ratio conditions, we found that ‘minis’ showed the best performance with the mean Area Under the (v-ROC) Curve (AUC) value of 0.960 ± 0.002 (95% confidence interval here onwards; Figure 5C). A paired samples t-test comparing to MiniAnalysis which had a mean AUC value of 0.937 ± 0.0022 gave t(13) = 12.4 and p = 1.37×10^-8^. Clampfit showed the poorest overall performance with the mean AUC value of 0.706 ± 0.0043, t(13) = 33.1, and p = 6.14×10^-14^ when compared to the MiniAnalysis mean AUC value in a paired samples t-test. These differences in detection performance were further corroborated by the d’ measure averaged across all signal-to-noise ratio conditions (Figure 5D). A paired samples t-test comparing mean d’ values from ‘minis’ (2.45 ± 0.045) versus MiniAnalysis (2.11 ± 0.04) gave t(13) = 14.0 and p = 3.24×10^-9^. The same test comparing the mean d’ from MiniAnalysis and Clampfit (1.23 ± 0.024) gave t(13) = 13.3 and p = 6.07×10^-9^. In summary, these findings clearly demonstrated that our ‘minis’ detection algorithm outperformed the other two algorithms, for ∼0.3 mV simulated minis added to real recording noise, at incidence rates above 10 minis/s, while Clampfit consistently showed the poorest detection performance across the entire incidence rate and signal-to-noise range examined, for these mPSP shapes and templates.

#### Detection under a wide range of incidence rate conditions: common errors

Both ‘minis’ and MiniAnalysis were fairly good at detecting moderately-sized (0.3 ± s.d. 0.05 mV) smEPSPs added to our noise-alone recordings. Yet all three algorithms, including ‘minis’, missed a sizeable proportion. Figure 6 shows examples of common mistakes. One type of mistake was missing simulated events that occurred very close to other events (Figures 6A, B, C, D, E, F, and G).

**Figure 6:**
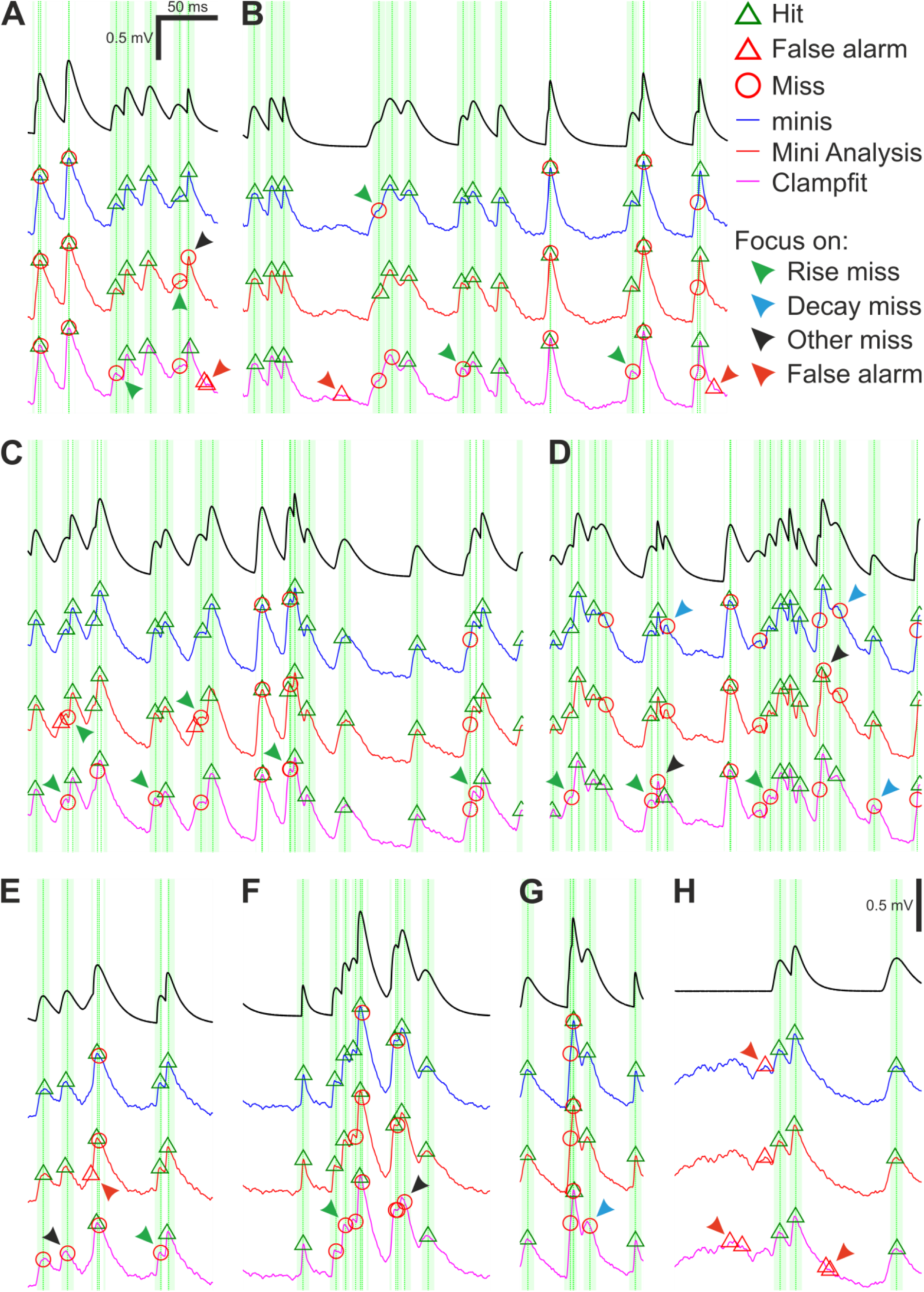
Moderately-sized (∼0.3 mV) smEPSPs (40 minis/s): detection errors made by the three algorithms. (A-H) Examples of smEPSP detection errors commonly committed by the three algorithms. Traces in blue, red, and magenta represent detection performance for ‘minis’, MiniAnalysis, and Clampfit, respectively. The top black trace shows actual smEPSPs. Vertical green dotted lines mark actual individual (i.e., non-summated) smEPSP peak locations with the fainter shaded colour demarcating a 10-ms window for accepting the detection of an mEPSP. Arrowheads indicate common detection errors made by the three algorithms. (Details in text).

Unsurprisingly, none of the three algorithms could deal with events that overlapped nearly perfectly, always lumping them together. Detecting peaks in the data using thresholds or fitting templates to the data cannot offer a way of identifying overlaps. Another common error type was missing events with peaks on the rising phase of another mEPSP (Figures 6A, B, C, D, E, and F; green arrowheads).

Unlike perfect or near-perfect overlaps, these events had a clear but brief decay period that was quickly upended by the rising phase of the next mini. Because these decays were brief, they were often lumped with a subsequent mEPSP or they could not be adequately fit with a template by Clampfit and, therefore, were discarded. The ‘minis’ program (but not MiniAnalysis) does offer a way to minimise these errors by reducing the peak integration period, but this comes at the expense of noise fluctuations on the rising phase increasingly passing as postsynaptic potentials (a type of false positive; Figure 6H, red arrowheads). An effective solution to this problem would, therefore, require isolating and measuring the noise component in whole-cell patch clamp recordings – which currently is not implemented in either MiniAnalysis or Clampfit (but is in ‘minis’).

An even more common error type was to miss (smaller) mEPSPs that occurred on the decay phase of another, often larger, earlier mEPSP (Figures 6D and G; blue arrowheads). The amplitude of mEPSPs occurring on decays is effectively reduced (as the peak drops relative to the baseline, due to the underlying decay of V_m_) and the steeper the decay, the more the amplitude is reduced. The amplitude of such a reduced mini may fall below the detection threshold and/or shape distortions may prevent fitting a template. As a result, such mEPSPs were occasionally discarded (this turns out to be a significant problem with real minis recorded from cortical pyramidal neurons, which exhibit significant temporal summation, because of their relatively high incidence rates and slow decays).

There is no easy way of correcting these errors, as it would (for example) require estimating the underlying decaying potential and subtracting it from the overall V_m_ or estimating the fall in the peak and adding it back to the amplitude before deciding whether to accept that candidate putative mini. mEPSPs that neither occurred during the rise nor decay phases were also occasionally missed (Figures 6A, D, E, and F; black arrowheads). This could have happened due to a failure to fit a template (Clampfit) or due to the amplitude being reduced below threshold by *downward* background noise fluctuations. In recordings of real mEPSPs, these errors would mainly affect small amplitude mEPSPs. Again, an effective solution to this problem would require isolating and measuring the noise component in whole-cell patch clamp recordings similarly to previously discussed errors occurring on the V_m_ rise phase.

A whole other class of detection errors appeared in the form of false alarms (false positives) occurring on the rising (Figure 6E; red arrowhead) or decaying phases (Figures 6A, B, and H; red arrowheads) of underlying mEPSPs or in the absence of any other minis in the vicinity (Figures 6B and H; red arrowheads). These mistakes were often caused by *upward* background noise fluctuations passing as real minis and might have been ‘enabled’ by low amplitude detection thresholds or an inability to discern shapes of real minis from mini-like noise. Just like the previously discussed error type, a solution would require isolating and measuring properties of the noise component. In addition, there were errors that uniquely affected only one of the algorithms, like the misplacement of detected events (peaks) by MiniAnalysis (Figure 6C; green arrowheads).

We saw that distinct detection errors could be made depending on how close they were to neighbouring mEPSPs and whether they occurred on the rise or decay phases of other nearby mEPSPs or in the absence of any other mEPSPs in the background. All three algorithms occasionally committed these errors, but their incidence was uneven.

#### Detection under a wide range of incidence rate conditions: the membrane potential ‘slope trend’ effect

In order to quantify the detection performance in different circumstances, for the same ∼0.3 mV smEPSPs, we evaluated the probability of making errors in relation to the *proximity* to neighbouring minis and the phase (*trend*) of the background V_m_: rising or decaying. We looked at TPR as a function of time interval to the nearest-neighbour mini (Figure 7A) taken together for all minis incidence rate conditions used above (but excluding conditions where noise was scaled): i.e., 640, 320, 160, 80, 40, 20, 10, 5, and 2.5 minis/s conditions (9 in total). We found that detection was superior for our ‘minis’ program across the entire inter-mini interval range. MiniAnalysis came second, with Clampfit consistently worst. These differences in detection performance were also reflected in the *cumulative* TPR (Figure 7D). As for the FPR (Figure 7B), ‘minis’ was superior at detecting ∼0.3 mV smEPSPs that were within 10 ms of the nearest neighbour (peak-to-peak). This performance edge over MiniAnalysis was not maintained for larger inter-mini peak intervals, while Clampfit again showed consistently the worst performance across the entire interval range. These performance differences were also reflected in the *cumulative* FPR (Figure 7E). Taken together, true and false positive rates allowed as to calculate a combined performance indicator, sensitivity index d’, as a function of time to the nearest neighbour (Figure 7C). The combined measure indicated that ‘minis’ had the superior detection performance within 10 ms of the nearest neighbour while Clampfit showed the worst performance across the entire inter-mini interval range.

**Figure 7:**
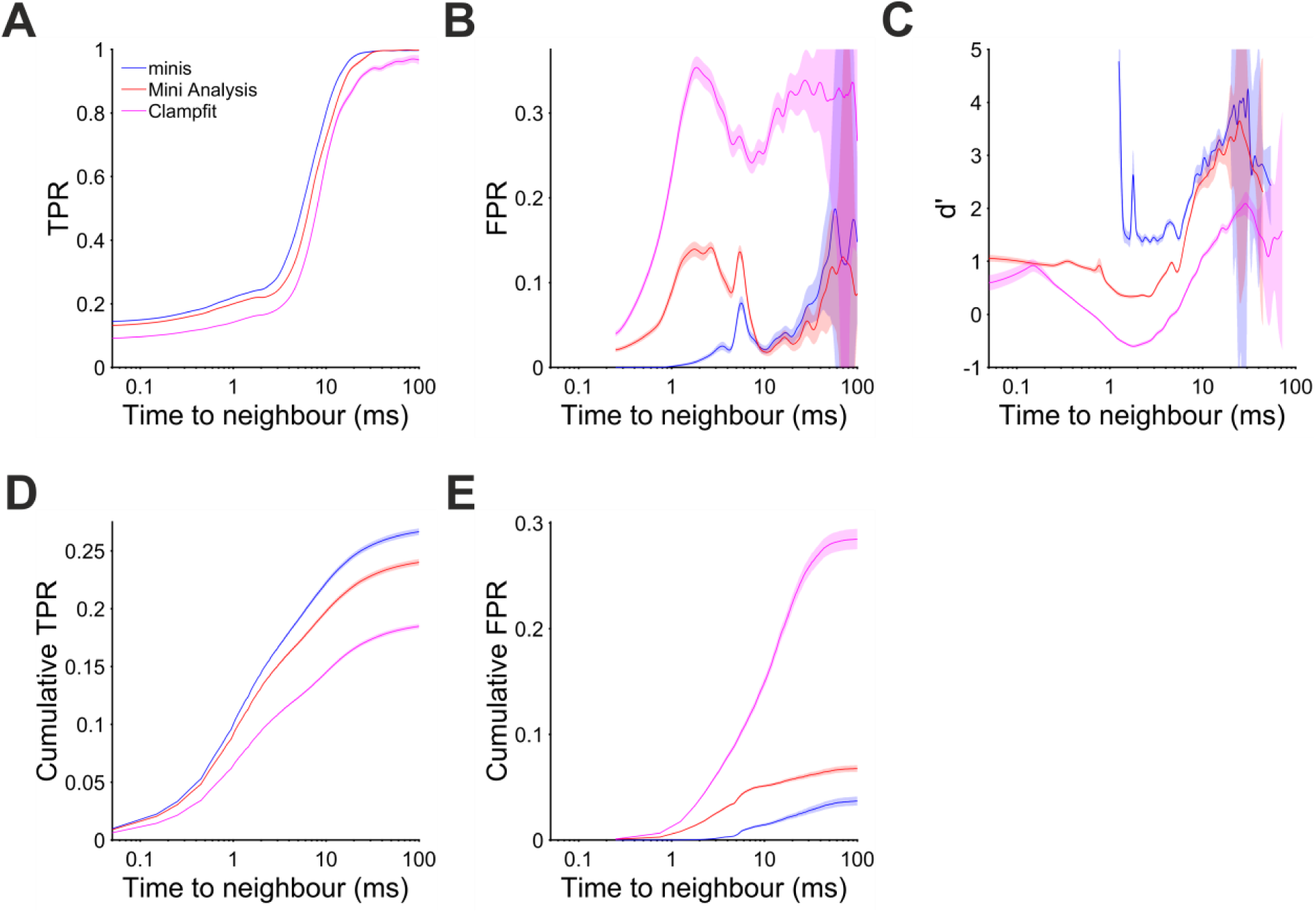
Performance when detecting moderately-sized (∼0.3 mV) smEPSPs as a function of time to the nearest neighbour (mean inter-mini peak-to-peak interval). Pale shaded colours indicate 95% confidence intervals. (A) True positive rate (TPR). (B) False positive rate (FPR). (C) Sensitivity index d’ (undefined but arbitrarily high for TPR > 0 and FPR = 0, e.g. for inter-mini intervals below 1 ms, for ‘minis’ algorithm). (D) Cumulative true positive rate (TPR) (cumulative true positives normalised by the total unbanded true positive count and scaled by the overall unbanded TPR; see text for details). (E) Cumulative FPR (cumulative false positives normalised by the total unbanded false positive count and scaled by the overall unbanded FPR).

During the V_m_ rise phase performance differences were unequivocal. There were very few events and, therefore, positive detections when the background V_m_ was rising at 200 μV/s or faster (Figure 8D). Most of the detected events coinciding with the rise phase actually occurred when V_m_ was changing at the rate of 10 to 100 μV/s. In this range ‘minis’ showed the best detection performance with d’ between 1 and 2 (Figure 8C). MiniAnalysis was the second best, while Clampfit showed the worst performance. This order of performance was maintained for rates below 10 μV/s with overall performance improving for all algorithms. The performance edge of ‘minis’ over the other algorithms was primarily due to low FPR (Figure 8B). The TPR of ‘minis’ was also superior to the other two algorithms but, in terms of sensitivity, MiniAnalysis was not far behind (Figure 8A). In the range between 15 and 35 μV/s ‘minis’ actually fell slightly behind MiniAnalysis. The cumulative rates clearly supported the observation that ‘minis’ was superior to the other two algorithms with MiniAnalysis coming second and Clampfit having the worst performance (Figures 8D and E).

**Figure 8:**
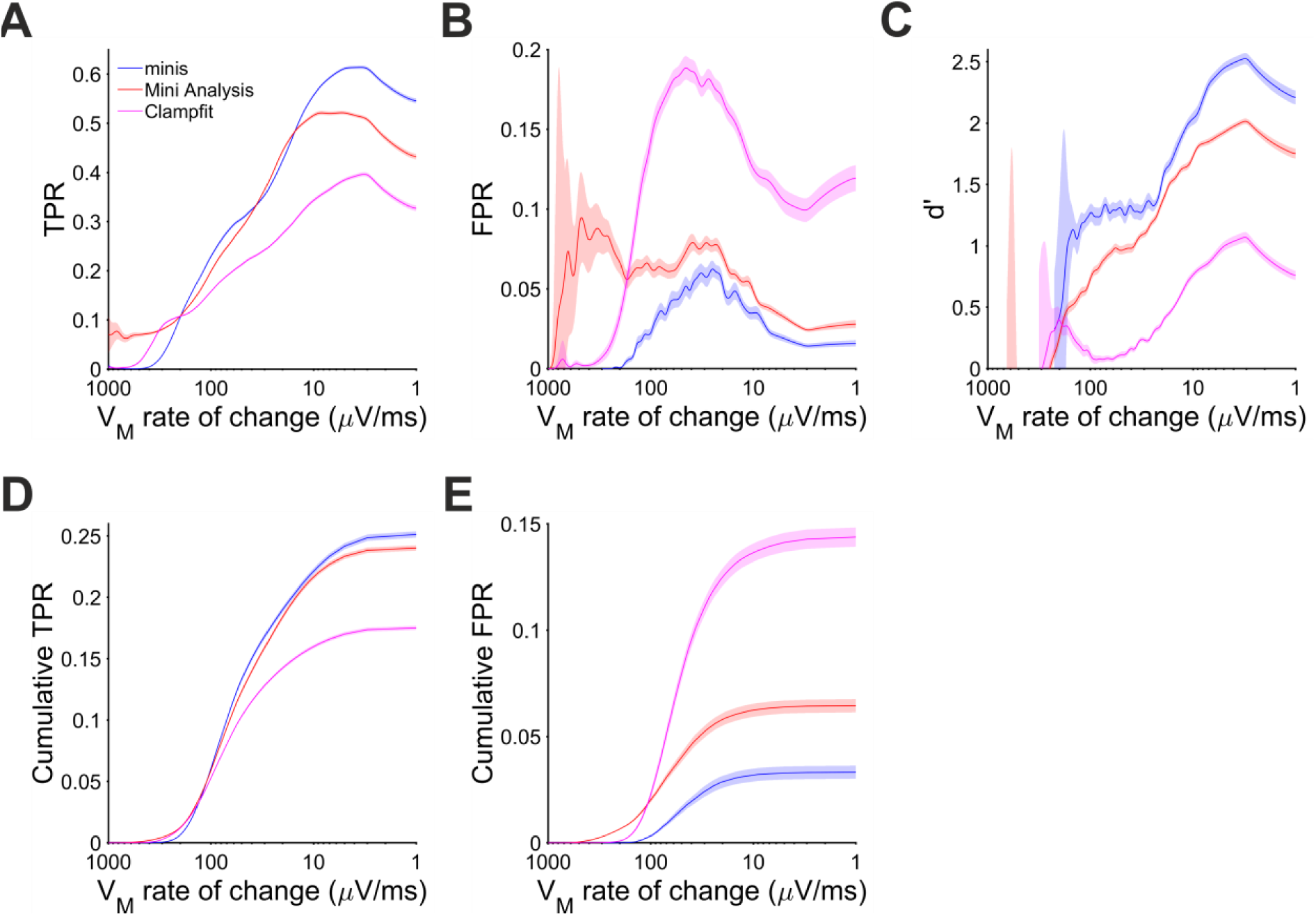
Performance when detecting moderately-sized (∼0.3 mV) smEPSPs on a rising trend V_m_ as a function of the V_m_ rate of change. Paler shaded colours indicate 95% confidence intervals. (A) True positive rate (TPR). (B) False positive rate (FPR). (C) Sensitivity index d’. (D) Cumulative TPR. (E) Cumulative FPR.

The same behaviour of the three algorithms was also observed for detecting ∼0.3 mV smEPSPs occurring on the decay phase of the background V_m_ (caused by summation of other smEPSPs). Most of the detected simulated events were associated with rates of V_m_ change in the range of –100 and – 10 μV/s (Figure 9D). In this range and above, ‘minis’ demonstrated the best performance relative to the other two algorithms with d’ ranging between 1.5 and 2.5 (Figure 9C). MiniAnalysis again came second, while Clampfit showed very poor performance. Clampfit often had a d’ that was close to 0, meaning that its performance was often not different from chance level when background V_m_ was decaying. This was mainly due to the large FPR for this algorithm (Figure 9B); ‘minis’ had unequivocally the highest TPR and the lowest FPR (Figures 9A and B) with cumulative rates very clearly reflecting this (Figures 9D and E). The overall performance of the runner-up, MiniAnalysis, was substantially poorer.

**Figure 9:**
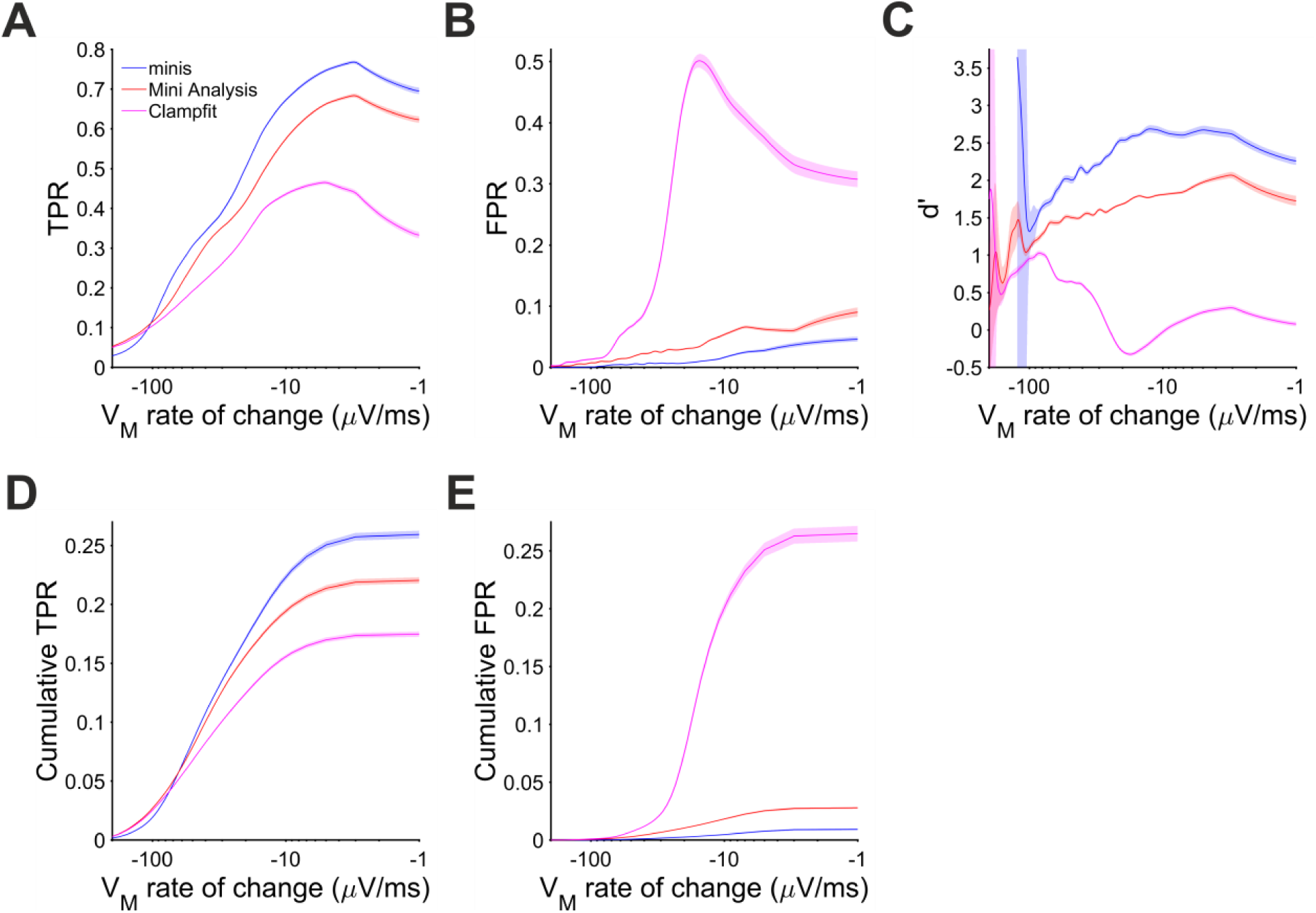
Performance when detecting moderately-sized (∼0.3 mV) smEPSPs on the membrane potential decaying trend phase as a function of the membrane potential rate of change. Paler shaded colours indicate 95% confidence intervals. (A) True positive rate (TPR) (B) False positive rate (FPR). (C) Sensitivity Index d’. (D) Cumulative TPR. (E) Cumulative FPR.

There are only 3 conditions with realistic incidence rates (20, 40, and 80 minis/s). Although, higher or lower incidence rates were useful in constructing the full v-ROC curve, it can be argued that they are artificial and experimental electrophysiologists may not be concerned with them. Therefore, we also examined the membrane potential ‘slope trend’ effect separately for the three realistic incidence rate conditions in the Supplementary Results section demonstrating similar findings to the overall results.

### Algorithm performance comparison for detecting simulated excitatory postsynaptic potentials with a wide set of discrete amplitudes, at realistic incidence rates

So far, we have only tested performance of the three algorithms when detecting moderately-sized (0.3 mV ± s.d. 0.05 mV) mEPSPs with a relatively narrow range of amplitudes. The existing evidence does not indicate that either mPSPs or mPSCs form a distinct amplitude distribution that can be separated from the amplitude distribution of noise fluctuations. Therefore, here we aimed to test the performance of detecting smEPSPs of a wide range of amplitudes, to evaluate the three algorithms in more realistic settings.

We used three incidence rate conditions: 27, 38, and 61 minis/s. Within each of these conditions smEPSPs were drawn pseudo-randomly from a uniform distribution of the following ten discrete amplitudes: 0.05, 0.1, 0.15, 0.2, 0.25, 0.3, 0.35, 0.4, 0.45, 0.5 mV (4 different simulations per noise trace per incidence rate conditions; see Methods for shapes). Just like in previous simulations, we positioned these smEPSPs pseudo-randomly over noise recordings and detected them using the three algorithms. The overall performance of these algorithms is shown in terms of true and false positive rates in Figure 10A and d’ in Figure 10B. A few things are prominent in these graphs. First of all, the detection performance of all algorithms was considerably worse than when detecting moderately-sized (∼0.3 mV) mEPSPs. Secondly, ‘minis’ again had the edge over two other algorithms, especially at higher minis’ incidence rates. Third, Clampfit’s performance was consistently the poorest, in our hands.

**Figure 10:**
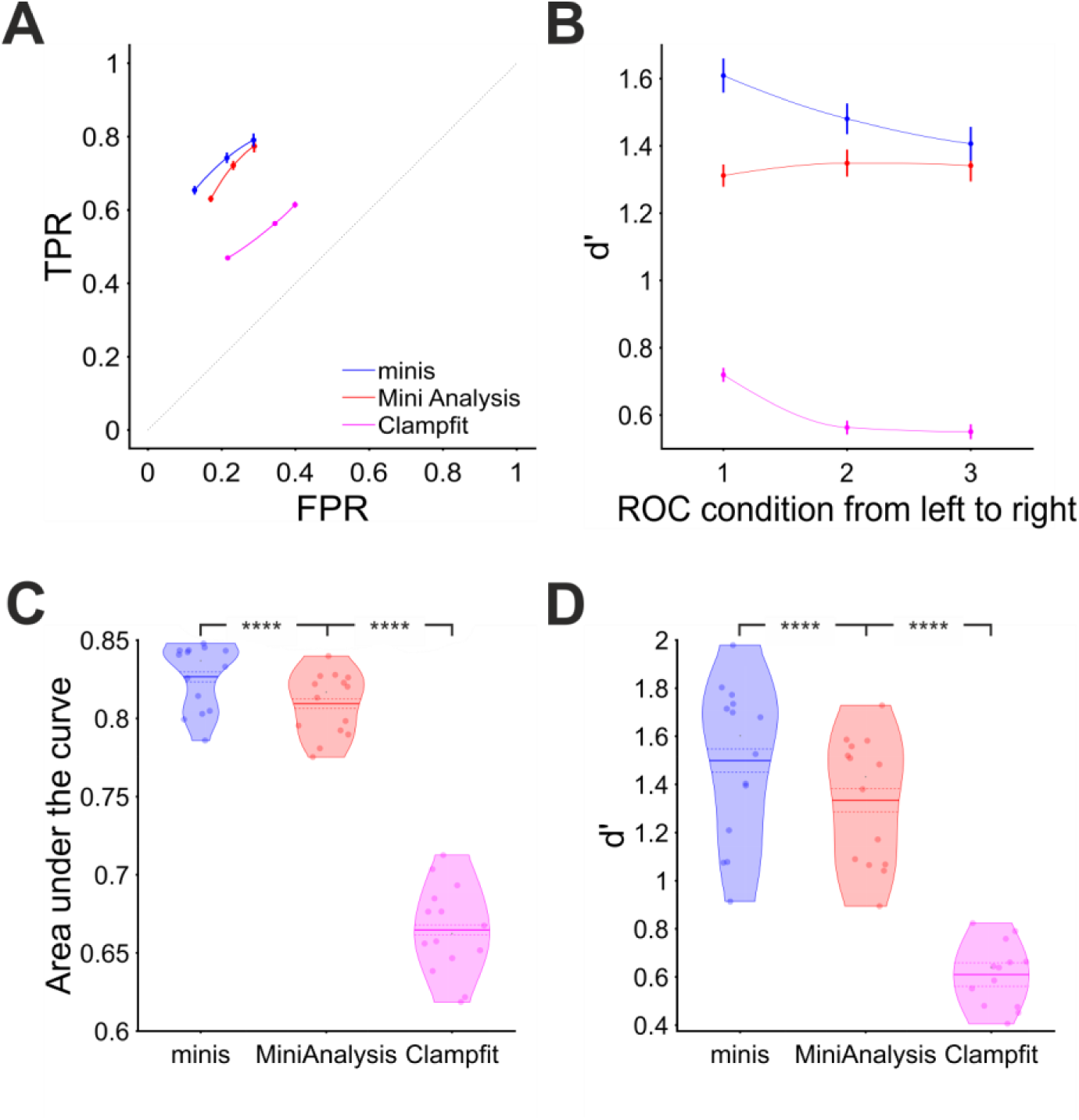
Overall performance for detecting smEPSPs of various (realistic) discrete amplitudes at ‘realistic’ minis’ rates (details in text). (A) Partial virtual “ROC” curve showing performance for detecting small-to-large-sized smEPSPs (0.05 to 0.5 mV in increments of 0.05 mV selected and added at random to a noise trace from each of the 14 cells) in terms of TPR (sensitivity) and FPR (1 – specificity) for all three algorithms. Moving from left to right different data points represent 3 different biologically realistic incidence rate conditions: 61, 38, and 27 minis/s. Vertical and horizontal bars indicate 95% confidence interval around the mean. The dotted diagonal line indicates chance performance. (B) Sensitivity index (d’) when detecting smEPSPs in the same 3 conditions as in (A) for all three algorithms. (C) Area under the virtual-ROC curve in (A) for all three algorithms. Individual data points represent individual recordings (n=14 cells), averaged over the three simulated minis incidence rates. The mean is marked by a solid line over the violin centre. The dashed line indicates the 95% confidence limits. **** indicates (highly) significantly different at p ≤ 10^-5^ level, paired t-test. (D) Sensitivity index d’ averaged across all three simulated minis incidence rates, for each cell (n=14), separately for each detection algorithm.

Performance was also assessed pooling the incidence rate conditions. We found ‘minis’ to have the best performance with the mean AUC value of 0.827 ± 0.003 (95% confidence interval from here on; Figure 10C). A paired samples t-test comparing to MiniAnalysis with the mean AUC value of 0.809 ± 0.003 gave p = 1.0×10^-6^. Clampfit showed the poorest performance, with the mean AUC value of 0.665 ± 0.004, p = 2.2×10^-10^ when compared to the MiniAnalysis mean AUC value in a paired samples t-test. These differences in detection performance were further corroborated by the d’ measure averaged across all incidence rate conditions (Figure 10D). The d’ discriminability index scores of ‘minis’ (1.5 ± 0.05) and MiniAnalysis (1.33 ± 0.04) were significantly different (p < 10^-5^), as were the d’ from MiniAnalysis and Clampfit (0.61 ± 0.02) (p < 4×10^-6^) The conclusion based on combined measures was in line with the assessment based on Individual incidence rate conditions.

Other measures like true positive and false positive rates as functions of the interval to the nearest neighbour and V_m_ rate of change during the rise or decay phases using more realistic settings pointed to a very similar conclusion to that made with the full virtual ROC curve. First, we found that ‘minis’ performed better than the other algorithms at detecting mEPSPs that were within 10 ms of other mEPSPs, and that at larger intervals, MiniAnalysis performed just as well (Figure 11).

**Figure 11:**
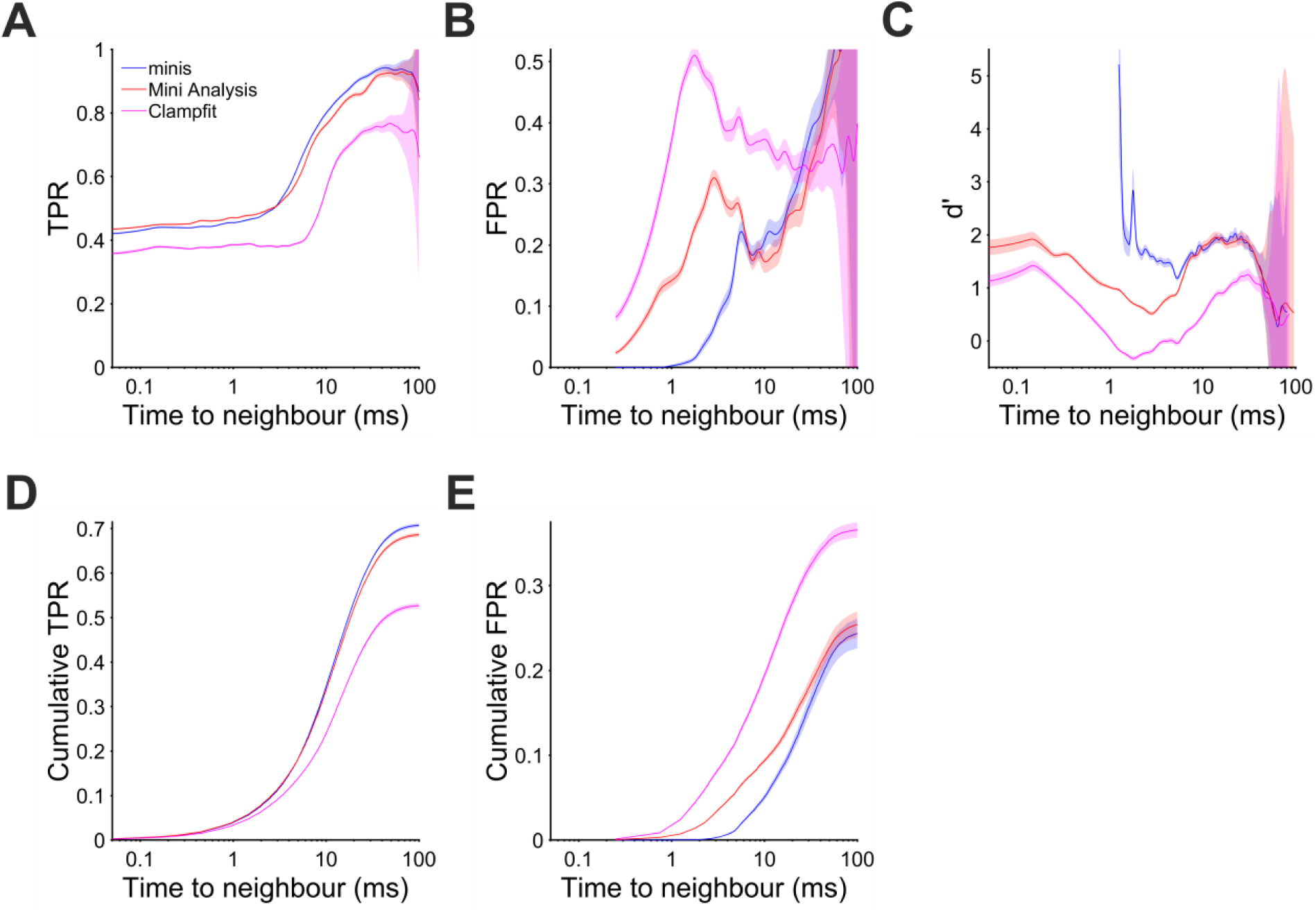
Performance for detecting smEPSPs of various (realistic) discrete amplitudes as a function of the time to the nearest neighbour. (A) True positive rate (TPR) when detecting smEPSPs of various discrete amplitudes (0.05 to 0.5 mV in increments of 0.05 mV) added at random to noise waveforms, as in the previous figure, as a function of the time to the nearest neighbour. Pale shaded colours indicate 95% confidence intervals. (B) False positive rate (FPR). (C) d’ (undefined but arbitrarily high for TPR > 0 and FPR = 0, e.g. for inter-mini intervals below 1 ms, for ‘minis’ algorithm). (D) Cumulative TPR. (E) Cumulative FPR.

Meanwhile, Clampfit performed consistently the worst across the range of inter-mini intervals whether in terms of d’ (Figure 11C) or true and false positive rates (Figures 11A, B, D, and E). When it came to V_m_ rate of change, ‘minis’ performed better than MiniAnalysis, which in turn performed better than Clampfit when detecting smEPSPs occurring during rise and decay phases of background simulated V_m_ (Figures 12 and 13). One difference compared to the full virtual ROC curve analysis was that the detection performance of ‘minis’ and MiniAnalysis overlapped in the range of 15-25 µV/s when detecting smEPSPs occurring on the rise phase of the background V_m_ (Figure 12C). This is largely caused by an increased FPR of ‘minis’ in this rate range (Figure 12B). The overall performance of all algorithms was better (Figures 12C and 13C) compared to the full virtual ROC curve analysis (Figures 8C and 9C), but still worse when compared to detecting moderately-sized (∼0.3 mV) smEPSPs with realistic incidence rates (Supplementary Figures 2C and 3C), as described in the previous subsection and in the Supplementary Results. In summary, ‘minis’ had a substantial advantage when detecting minis in close proximity to neighbouring minis, as well as during rising or falling background V_m_, with Clampfit severely underperforming.

**Figure 12:**
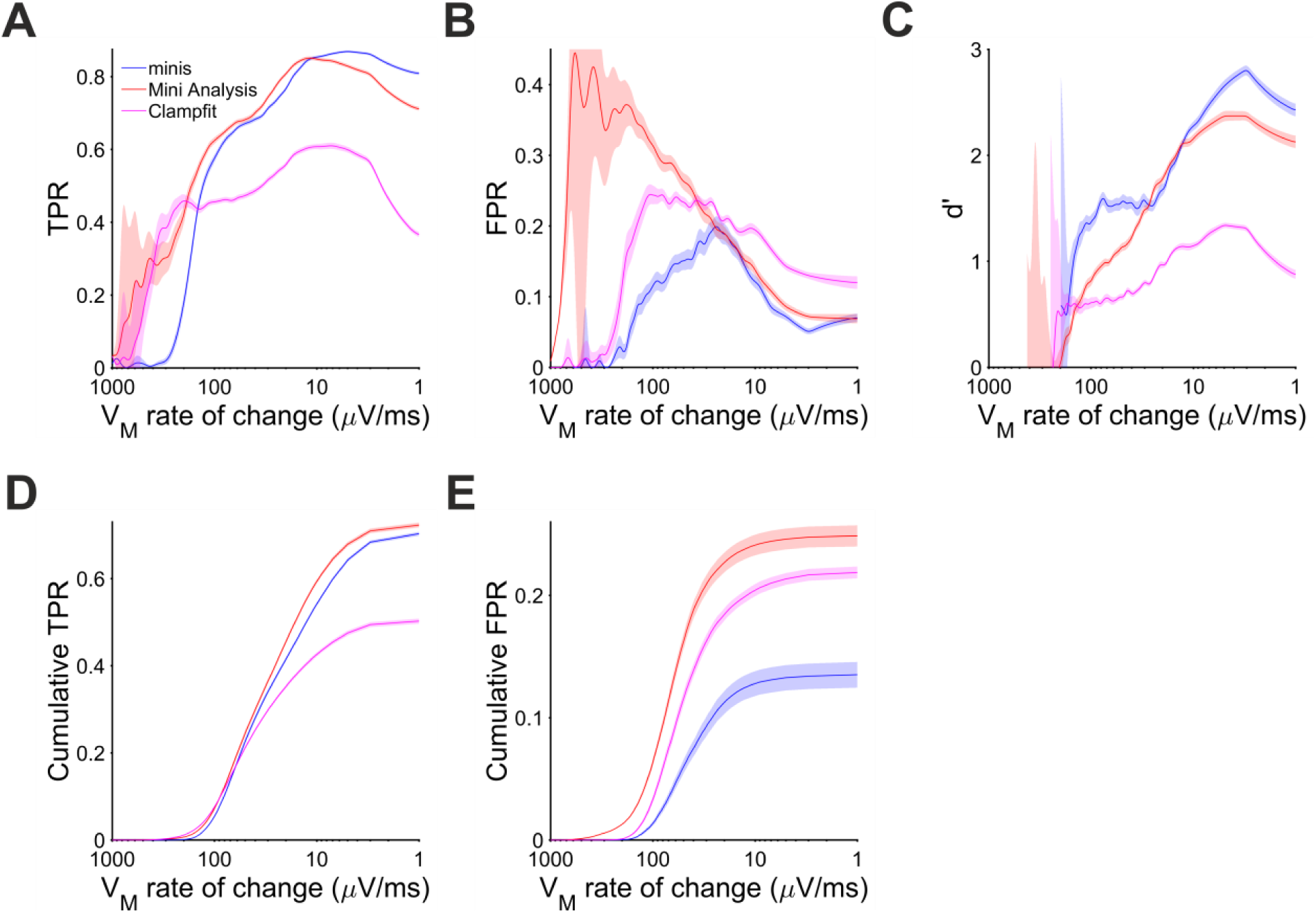
Performance for detecting smEPSPs of (realistic) discrete amplitudes on the membrane potential rise phase. (A) True positive rate (TPR) when detecting smEPSPs of various amplitudes (0.05 to 0.5 mV in increments of 0.05 mV, added at random to noise waveforms, as in the previous figure), on the V_m_ rise phase as a function of the V_m_ rate of change. Pale shaded colours indicate 95% confidence intervals. (B) False positive rate (FPR). (C) d’. (D) Cumulative TPR. (E) Cumulative FPR.

**Figure 13:**
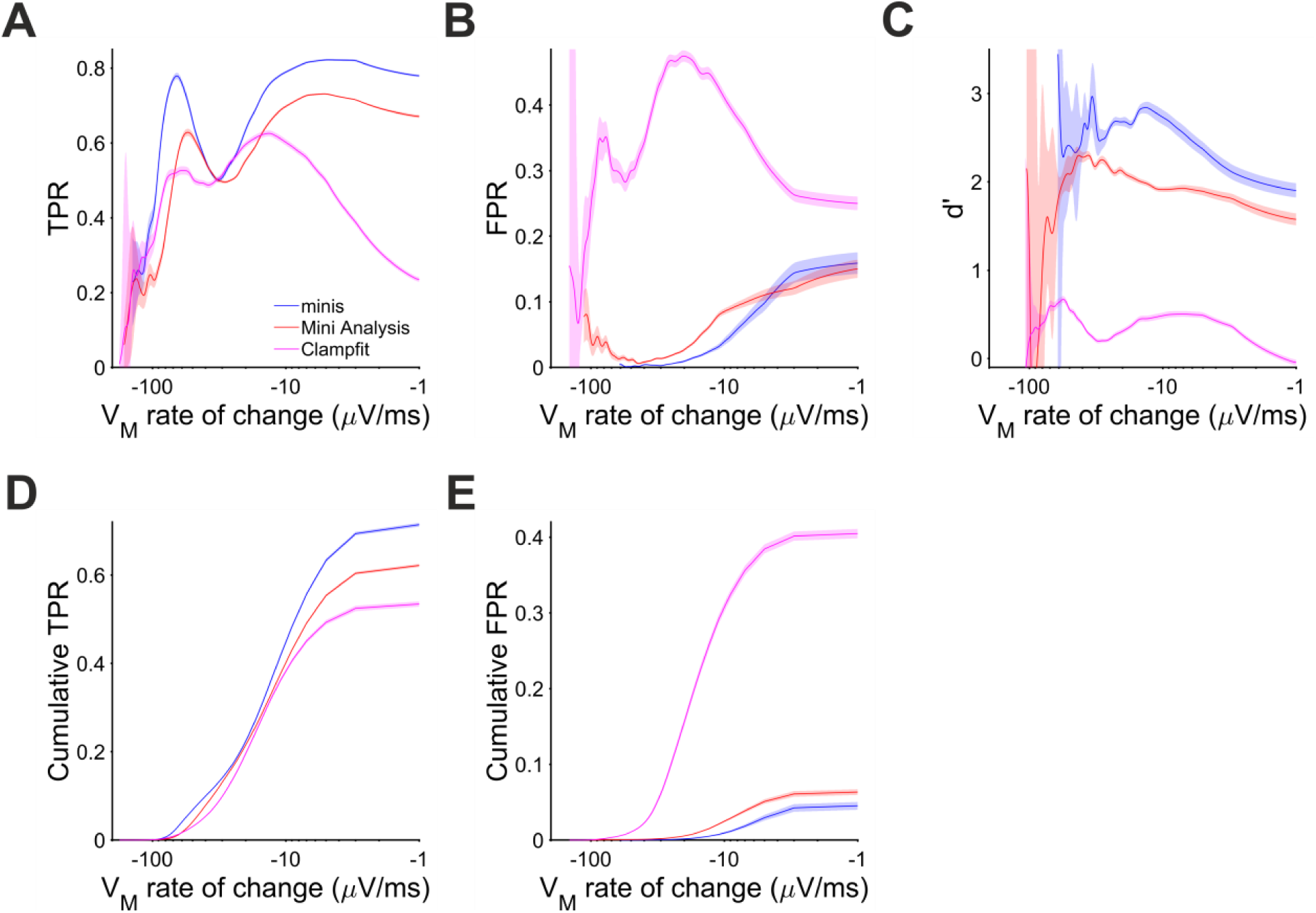
Performance for detecting smEPSPs of (realistic) discrete amplitudes on the membrane potential decay phase under realistic incidence rate conditions. (A) True positive rate (TPR) when detecting smEPSPs of various amplitudes (0.05 to 0.5 mV in increments of 0.05 mV, as in the previous figure) on the V_m_ decay phase, as a function of the V_m_ rate of change. Pale shaded colours indicate 95% confidence intervals. (B) false positive rate (FPR). (C) d’. (D) Cumulative TPR. (E) Cumulative FPR.

### Comparison of amplitudes, time course, and incidence rates of detected real and simulated excitatory postsynaptic potentials by different algorithms

In the final section we looked at how the three tested algorithms fare across all simulation conditions in terms of their minis’ amplitude, time course, and incidence rate estimates. We also carried out detection of real minis in the same recorded cells prior to blocking minis (‘noise with minis’ condition) and compared the same estimates for the three algorithms. A large number of estimates and inferential statistics was compiled into several tables providing a thorough performance overview.

Table 1 below shows the compiled mean amplitude estimates for simulated and real minis detected by the three algorithms while Supplementary Table 1 shows repeated samples t-test p-values for comparisons of mean amplitude values for the three different algorithms with respect to ground truth values. Mean amplitude values of simulated and real mPSPs detected by ‘minis’ were consistently lower than mean amplitude values of events detected by both MiniAnalysis and Clampfit and were closest to ground truth values. Detection of simulated events by the ‘minis’ algorithm in conditions with realistic smPSP incidence rates (i.e., Type 1 RI and Type 2 rows in the tables) were extremely close to the ground truth values (0.3 mV vs. 0.31 mV and 0.275 mV vs. 0.29 mV, respectively; Table 1). These observations further confirm that ‘minis’ detection algorithm is exceptionally good at picking up small PSPs. The tendency to overestimate amplitudes of detected smPSPs in high incidence conditions (i.e., Type 1 HI row in the tables) by all three algorithms reflects the increased probability of major overlap between simulated events in these conditions.

**Table 1:**
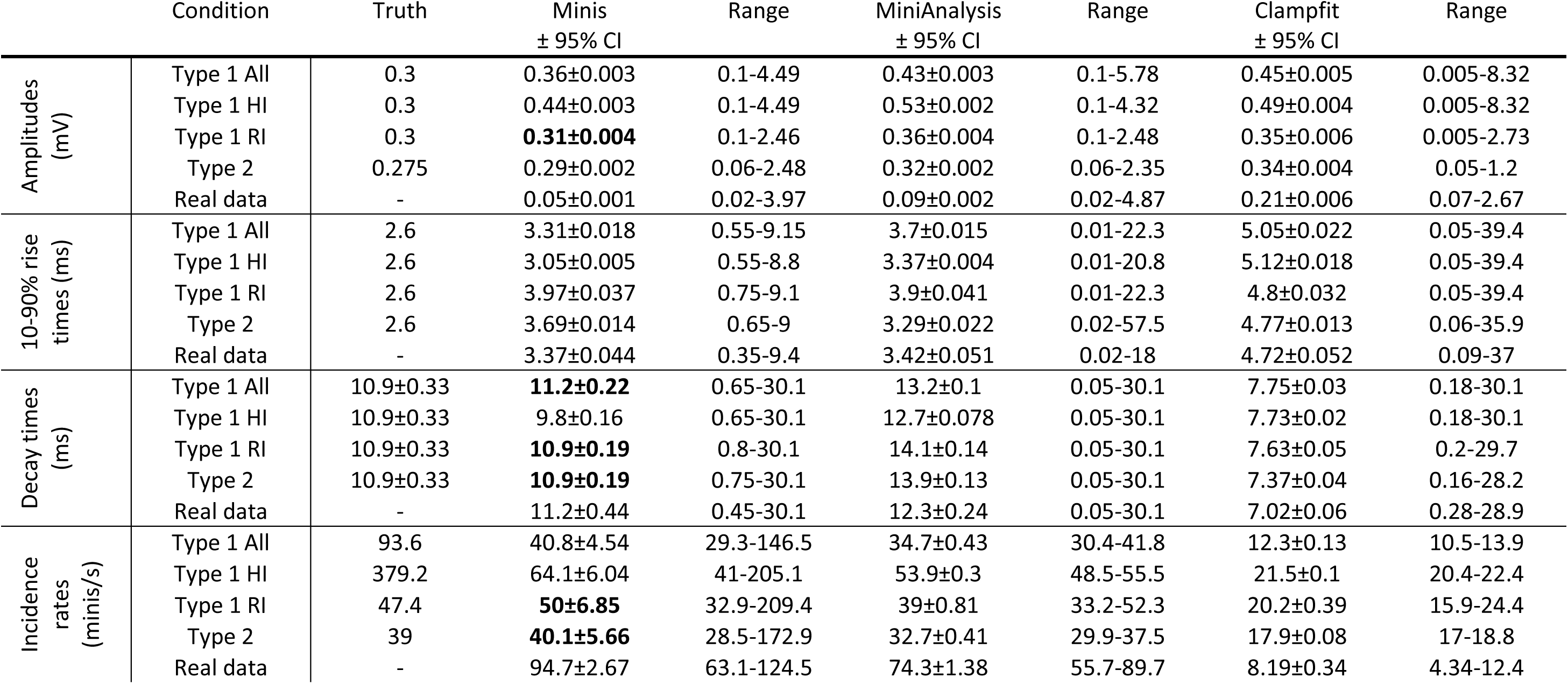
Mean amplitudes, 10-90% rise times, decay times, and incidence rates of detected simulated and real minis by the three different algorithms. Abbreviations: Confidence interval (CI), all conditions with 0.3 mV amplitude simulated minis used to construct the virtual ROC curve (Type 1 All), high incidence rate conditions (640, 320, and 160 minis/s) with 0.3 mV amplitude simulated minis used to construct the virtual ROC curve (Type 1 HI), realistic incidence rate conditions (80, 40, and 20 minis/s) with 0.3 mV amplitude simulated minis used to construct the virtual ROC curve (Type 1 RI), all realistic incidence rate conditions with varying amplitude simulated minis (Type 2). Statistically not significant comparisons to ground truth values are emphasised in bold (p-values are provided in Supplementary Table 1).

Table 1 below shows the compiled mean 10-90% rise time estimates for simulated and real minis detected by the three algorithms while Supplementary Table 1 shows repeated samples t-test p-values for comparisons of mean 10-90% rise time values for the three different algorithms with respect to ground truth values. All three algorithms tended to significantly overestimate the rise times of detected minis in all simulated conditions. This can be explained by the fact that noise fluctuations affect the baseline positioning by shifting it to the lowest trough some time prior to the real start of the event. The rise times of detected events are also more likely to be overlaps of more than a single event and, therefore, would tend to be longer if lumped together. This was especially true for events detected by Clampfit which had a pronounced tendency to lump smaller events into fewer bigger ones. Both MiniAnalysis and Clampfit had large range values. For example, MiniAnalysis had a range of 0.02-18 ms and Clampfit had a range of 0.09-37 ms when detecting real mPSPs, while ‘minis’ had a range of 0.35-9.4 ms when detecting the same type of events (Table 1). This observation is indicative of the fact that MiniAnalysis and Clampfit occasionally failed to accurately position the baseline of a detected event and failed to reject ‘wildly’ inaccurate estimates.

Table 1 below shows the compiled mean 1/e decay time estimates for simulated and real minis detected by the three algorithms while Supplementary Table 1 shows repeated samples t-test p-values for comparisons of mean decay time values for the three different algorithms with respect to ground truth values. Decay times of smPSPs detected by ‘minis’ were the closest to the ground truth values when compared to the other two algorithms. For simulated realistic incidence rate conditions (i.e., Type 1 RI and Type 2 rows in the tables) p-values were not significant for the ‘minis’ vs. the ground truth comparison (Table 1). Unfortunately, the same cannot be said of MiniAnalysis and pClamp which consistently over– and underestimated mean decay times, respectively (Table 1 and Supplementary Table 1). These differences between the algorithms may stem from differences in how decays are being estimated: Whether effective decays are measured or detected events are fitted with exponential decay functions. ‘Minis’ reduce these discrepancies by averaging the two methods to obtain a single estimate.

Table 1 below shows the compiled mean incidence rates for simulated and real minis detected by the three algorithms while Supplementary Table 1 shows repeated samples t-test p-values for comparisons of mean incidence rates for the three different algorithms with respect to ground truth. Values produced by ‘minis’ and MiniAnalysis were close to each other as well as ground truth values (albeit less so for MiniAnalysis) as indicated by corresponding p-values of the t-test statistic (Table 1 and Supplementary Table 1). Incidence rate values produced by Clampfit were considerably lower than the other two algorithms or the ground truth values indicating that Clampfit missed many events, as well as tended to lump events together. A distinct ability by ‘minis’ algorithm to pick up smaller events was further supported by a considerably larger incidence estimate for real data (comparable to the range of incidence rates reported in our companion article (Dervinis and Major, 2024)) when compared with the other two algorithms. Clampfit values for real data were smaller by almost an order of magnitude revealing a really poor performance by this algorithm.

## Discussion

Our analysis suggests that mini (mEPSC/P) incidence rates typically reported in neocortical slices in the CNS synaptic function literature may be underestimated, by up to an order of magnitude. Discrepancy of this size indicates that signal-to-noise ratio is a severe issue preventing accurate detection of minis irrespective of the detection method used: That is, both ‘template matching’ and ‘thresholded amplitude detection’ approaches are significantly flawed in distinguishing real minis from the background physiological noise. Having stated that, however, we also demonstrated that the two approaches are not equally flawed and that ‘thresholded amplitude detection’ performs reasonably well and is further improved by our novel mEPSC/P detection algorithm called ‘minis’.

We quantified the performance of ‘minis’ using standard measures from signal detection theory – namely, ROC (receiver operating characteristic) curves and sensitivity index d’, alongside measures of sensitivity – true positive rate (TPR) and false positive rate (FPR = 1 – specificity) – and demonstrated superior performance of our algorithm relative to the other two most popular algorithms in the field of minis research: MiniAnalysis and Clampfit. Our algorithm had a performance edge over the other two algorithms not only for detecting moderately-sized (∼0.3 mV) mEPSPs, but also across a wide range of realistic mEPSP amplitudes, shapes and incidence rates. Crucially, the latter included the higher minis’ incidence rates observed in neocortical slice experiments at body temperature, 13-80 minis/s, where temporal summation becomes an increasing challenge for detection. As a necessary step in building a reliable quantal size estimation method, we succeeded in developing an algorithm for detecting postsynaptic events that is transparent, systematically evaluated, and flexible.

Notwithstanding its good performance, the ‘minis’ algorithm has important limitations. First, the ‘minis’ algorithm, just like the other two detection algorithms, cannot by itself alone be used to estimate quantal sizes. Due to amplitude attenuation of spontaneous postsynaptic events and the need to use amplitude detection thresholds, the estimated amplitude will typically only be an upper limit on the actual population mean amplitude (unless a very low threshold is used in which case FPR would increase). We have also demonstrated that mini detection is impaired on rising and decaying trends of V_m_. The severity of this problem worsens with the rate of change of V_m_. The problem is exacerbated as the time between neighbouring minis gets shorter. Therefore, the mean amplitude estimate is not only affected by the presence of noise and by amplitude attenuation, but also by the presence of other miniature postsynaptic events. All the above limitations also apply to estimating the incidence rate of postsynaptic events. All the mentioned factors reduce event detectability and, therefore, result in an underestimated incidence rate. However, ‘detecting’ noise events as false minis can cause errors in the other direction if too low a detection threshold is set. Despite these limitations, our program ‘minis’ fares better on this front compared to the other two algorithms: ‘minis’ performance is also superior on rising and decaying trends in V_m_, and for closely-spaced events. To our knowledge, this is the first time these problems have been addressed, and their impact quantified in a systematic manner, and their implications discussed.

Other caveats concern the method for assessing the performance of detection algorithms. We did not use the full plausible range of realistic distributions for simulating mEPSPs in terms of their incidence rates, amplitudes, and their rise and decay times. However, the shapes of these distributions are not entirely known, and this is an empirical question that remains unanswered. Because of this reason, we refrained for trying to address this question here and avoided making empirically dubious assumptions. Instead, we explored a range of scenarios where simulation incidence rates ranged from more to less realistic. We explored the effects of varying signal/noise ratios. We believe that the large size of the parameter space we have explored does allow us to conclude that the three algorithms differ systematically in their performance, with our algorithm – ‘minis’ – being superior to the other two (in our hands at least).

We have not constructed ROC curves in the usual fashion by varying a detection threshold of each algorithm. Unfortunately, we were limited by the nature of the algorithms, specifically Clampfit which does not use a pure amplitude detection threshold but uses convolution of the data trace with each of the templates. However, our manipulations of varying the incidence rate of simulated events, as well as varying the amplitude of noise, allowed us to mimic effects of varying the detection threshold. We are also aware of one particular issue regarding the classification of correct rejections (true negatives). One could argue that a proper definition of correct rejections would require treating every data sample point as a potential noise event that could either be falsely detected as a signal event or correctly rejected as a noise event. Unfortunately, such a definition would result in large numbers of correct rejections (true negatives) swamping the number of other detection measures (true positives, false alarms (false positives), and misses (false negatives)) and, therefore, would make the detection performance analysis essentially meaningless. Alternatively, treating every 10 ms noise window as a basis for correct rejection would still produce a relatively large number of correct rejections, as would similar, more complicated schemes. There is also no reason why a single noise event should be limited to a single 10 ms window. Therefore, we thought that a reasonably practical definition of correct rejections as rejected prominent noise fluctuations fared better than the alternatives. Having explicit, transparent, and reproducible, methods to benchmark the performance is better than relying on subjective examination of recording traces.

Finally, we would like to address the question of whether we used optimal detection settings for each algorithm. MiniAnalysis settings were similar to those of ‘minis’. The two algorithms are somewhat similar and chosen settings were observed to give the best performance for both algorithms when detecting simulated mPSPs, after considerable exploration. In case there is any doubt about our method, we have made our recorded and simulated data and Matlab analysis code publicly available. With regards to using Clampfit to analyse simulated data, we used 9 templates (the maximum allowed) constructed in a typical fashion by averaging a number of waveforms with similar shapes but separating qualitatively different shapes. These templates have also been made available publicly. We tested different detection regimes with templates based on both smoothed or unsmoothed traces and comparing detection of smoothed and unsmoothed recording traces and chose the best regime. However, we did not construct new templates for every new recording or every new simulation condition, because this is extremely time-consuming and is not an efficient way to conduct this type of analysis, although Clampfit detection performance might have improved marginally. Clampfit might perform better if the number of templates was not limited to 9, and if a wide range of inbuilt templates was available. These issues are, however, outside the scope of this study. Given these considerations, we are confident in the validity of our approach.

We defer the discussion of possible reasons why there is such a big discrepancy between reported and expected minis’ incidence rates in the synaptic physiology literature and how they can be effectively addressed to our companion article (Dervinis and Major, 2024). It is worth noting, however, that ‘minis’ or another similar algorithm can be used to objectively measure the properties of pure noise fluctuations (with minis blocked). This would allow one, in principle, to separate ‘signal’ (minis) from noise components in traces of minis recorded in the presence of noise. Noise events could potentially be ‘removed’ from the combined signal and noise event distribution by direct histogram subtraction, although in practice it turns out one needs to compensate for the decay of the underlying waveform, following minis, and noise peaks coinciding with minis peaks: noise and pure noise-free minis histograms don’t simply summate (Dervinis and Major, 2024).

Alternatively, noise traces can be used to more accurately ‘reverse-engineer’ the signal distribution via simulations, by iteratively adjusting the shapes, amplitudes, and incidence rates of simulated minis, and adding these to real noise, under the control of an optimisation algorithm to match up the distributions of *simulated* minis plus real noise to *real* minis plus real noise. The companion article (Dervinis and Major, 2024) explores this and presents a quantal analysis method based on the ‘minis’ spontaneous postsynaptic event detection algorithm.

## Acknowledgements

Cardiff University (GM, MD), The Medical Research Council of the UK (MD 19 PhD). Neural activity simulations were performed on the Cardiff School of Biosciences’ Biocomputing 20 Hub HPC/Cloud infrastructure.

## Abbreviations

aCSF: artificial cerebrospinal fluid
AMPA: α-amino-3-hydroxy-5-methyl-4-isoxazolepropionic acid
AP: action potential
API: application programming interface
AUC: area under the curve
CNS: central nervous system
CPP: (*RS*)-3-(2-carboxypiperazin-4-yl)-propyl-1-phosphonic acid
d’: sensitivity index ‘d-prime’ (defined in Methods)
EPSC: excitatory postsynaptic current
EPSP: excitatory postsynaptic potential
FNR: false negative rate (‘miss’ rate)
FPR: false positive rate (‘false alarm’ rate)
GABA: gamma-aminobutyric acid
IPSC: inhibitory post-synaptic current
IPSP: inhibitory post-synaptic potential
mEPSC: miniature excitatory post-synaptic current
mEPSP: miniature excitatory post-synaptic potential
mIPSC: miniature inhibitory post-synaptic current
mIPSP: miniature inhibitory post-synaptic potential
mPSC: miniature post-synaptic current
mPSP: miniature post-synaptic potential
NBQX: 2,3-dioxo-6-nitro-1,2,3,4-tetrahydrobenzo[*f*]quinoxaline-7-sulfonamide
NMDA: N-methyl-D-aspartate
ROC: receiver operator characteristic
sEPSC: spontaneous excitatory post-synaptic current
sEPSP: spontaneous excitatory post-synaptic potential
sIPSC: spontaneous inhibitory post-synaptic current
sIPSP: spontaneous inhibitory post-synaptic potential
smEPSC: simulated mEPSC
smEPSP: simulated mEPSP
smPSC: simulated mPSC
smPSP: simulated mPSP
sPSC: spontaneous post-synaptic current
sPSP: spontaneous post-synaptic potential
TNR: true negative rate
TPR: true positive rate (‘hit’ rate, sensitivity)
TTX: tetrodotoxin
V_m_: membrane potential
v-ROC: virtual ROC (ROC-like plot)

## Supplementary Results

### Research literature measuring minis in the central nervous system: Extended report

There were 107 studies in total that matched our criteria. All of these studies used voltage clamp as the electrophysiological method for recording spontaneous and miniature postsynaptic events. We surveyed the software they used to detect these events. The biggest group, ∼48%, reported using MiniAnalysis. 26% reported using Clampfit, while 13% used a custom algorithm. The remaining 13% did not specify. We compared the performance of the two most popular detection algorithms, MiniAnalysis and Clampfit, both commercially available, with our new algorithm ‘minis’.

35 studies (ca. 32%) reported using a current amplitude detection threshold of which 19 studies specified actual numeric value used (–6.8 ± s.d. 0.44 pA, range of –3 to –20 pA). It is likely that a higher fraction of studies made use of amplitude thresholds as only two explicitly reported not using any detection thresholds. Most of the studies did not give enough details (65% of all studies) to judge one way or the other. However, studies that used MiniAnalysis for detection must have used a detection threshold, as this software requires it.

Out of all the studies, 18 carried out research in neocortical pyramidal cells looking at mPSCs only (spontaneous PSCs excluded). 12 studies reported a mean mEPSC amplitude value of –11.74 ± s.d. 0.77 pA (range of –6.0 to –20.9 pA). 13 studies reported a mean incidence rate value of 3.6 ± s.d. 0.34 per second (range of 1.0 to 8.0 per second).

The total putative excitatory synapse count on a single cortical pyramidal cell can number in the tens of thousands (Eyal et al., 2018b). Layer 2/3 pyramidal cells in rats are thought to have 5,000 to 30,000 putative excitatory synapses (Larkman, 1991; Eyal et al., 2018b). Large-thick-trunk/tufted layer 5 pyramidal cells would have an even larger number of contacts (Larkman, 1991). Therefore, we can roughly assume that for most cortical pyramidal cells the number of putative excitatory synapses ranges between 5,000 and 50,000. Conservatively, we could assume that only a minority, roughly 40%, of them are functional (‘non-silent’) synapses with active zones and post-synaptic densities containing AMPA receptors (Holler et al., 2021; Motta et al., 2019). Given that a spontaneous vesicle release at a single release site has been estimated to occur with an approximate incidence rate of 0.0021 per second (Murthy and Stevens, 1999) and given that there may be at least 2.7 release sites per cortical synapse (Holler et al., 2021), the lower bound incidence rate estimate of such events within a single cortical pyramidal neuron might be expected to range somewhere between ca. 10 minis/s and 100 minis/s (5,000*0.4*0.0021*2.7– 50,000*0.4*0.0021*2.7). Incidence rates values reported in the recent literature are below this predicted range by an order of magnitude, suggesting a significant fraction of smaller minis may commonly be ‘missed’ under some recording conditions. This problem has been observed directly (Nevian et al., 2007), using paired dendritic and somatic whole-cell patch recordings, albeit using voltage recording rather than voltage clamp, recording from large layer 5 neocortical pyramidal neurons.

### Detection under realistic incidence rate conditions (20, 40, 80 minis/s)

Data points used to construct the full v-ROC curve include incidence rate and noise conditions that were not necessarily realistic. Based on our calculations, the range of mEPSP incidence rates one might commonly expect during real recordings from neocortical pyramidal neurons is likely to be between 10 and 100 minis/s (see above and (Dervinis and Major, 2024)); In Figures 5A and B, 20, 40 and 80 minis/s (from right to left) are highlighted by open circles. Just as across the entire ROC curve, the performance of ‘minis’ was superior in this incidence rate range compared to the other two algorithms with MiniAnalysis coming second, and Clampfit being worst, by a substantial margin.

Other measures like true positive and false positive rates as functions of the time to the nearest neighbour and V_m_ rate of change during the rise or decay phases pointed to a very similar conclusion made regarding the full ROC curve. First, we found that ‘minis’ performed better than other algorithms at detecting mEPSPs that were within 10 ms of other mEPSPs and that at longer delays MiniAnalysis performed just as well (Supplementary Figure 1). Meanwhile, Clampfit performed consistently worst across the range of inter-mini intervals, whether measured by d’ (Supplementary Figure 1C) or true positive and false positive rates (Supplementary Figures 1A, B, D, and E). When it came to V_m_ rate of change, ‘minis’ consistently showed the best performance at detecting smEPSPs occurring with the background V_m_ rate of change ranging between –40 and 100 µV/s (Supplementary Figures 2 and 3). This is the range where the vast majority of simulated events occurred (Supplementary Figures 2D and 3D). In terms of individual TPR and FPR (Supplementary Figures 2A and B and 3A and B) and cumulative rates (Supplementary Figures 2D and E and 3D and E), the conclusions are much the same as those made regarding the full virtual ROC curve. Namely, that ‘minis’ had the largest TPR and the smallest FPR with an exception being the TPR in relation to rising background (simulated) membrane potential trends (Supplementary Figures 2A and D), with there being a range (45-100 µV/s) where MiniAnalysis had a slightly higher TPR, but not overall performance measured as d’ (Supplementary Figure 2C). Similar to conditions used to construct the full ROC curve, Clampfit showed consistently the worst performance by all measures. The performance of all three algorithms was better with realistic smEPSP incidence rates (Supplementary Figures 2C and 3C) than over the full ‘virtual’ ROC curve (Figures 8C and 9C). In summary, evaluation of detection of mEPSPs simulated at realistic incidence rates supported conclusions made regarding detection performance under a much broader range of simulation incidence rates.

## Supplementary Figures

**Supplementary Figure 1:**
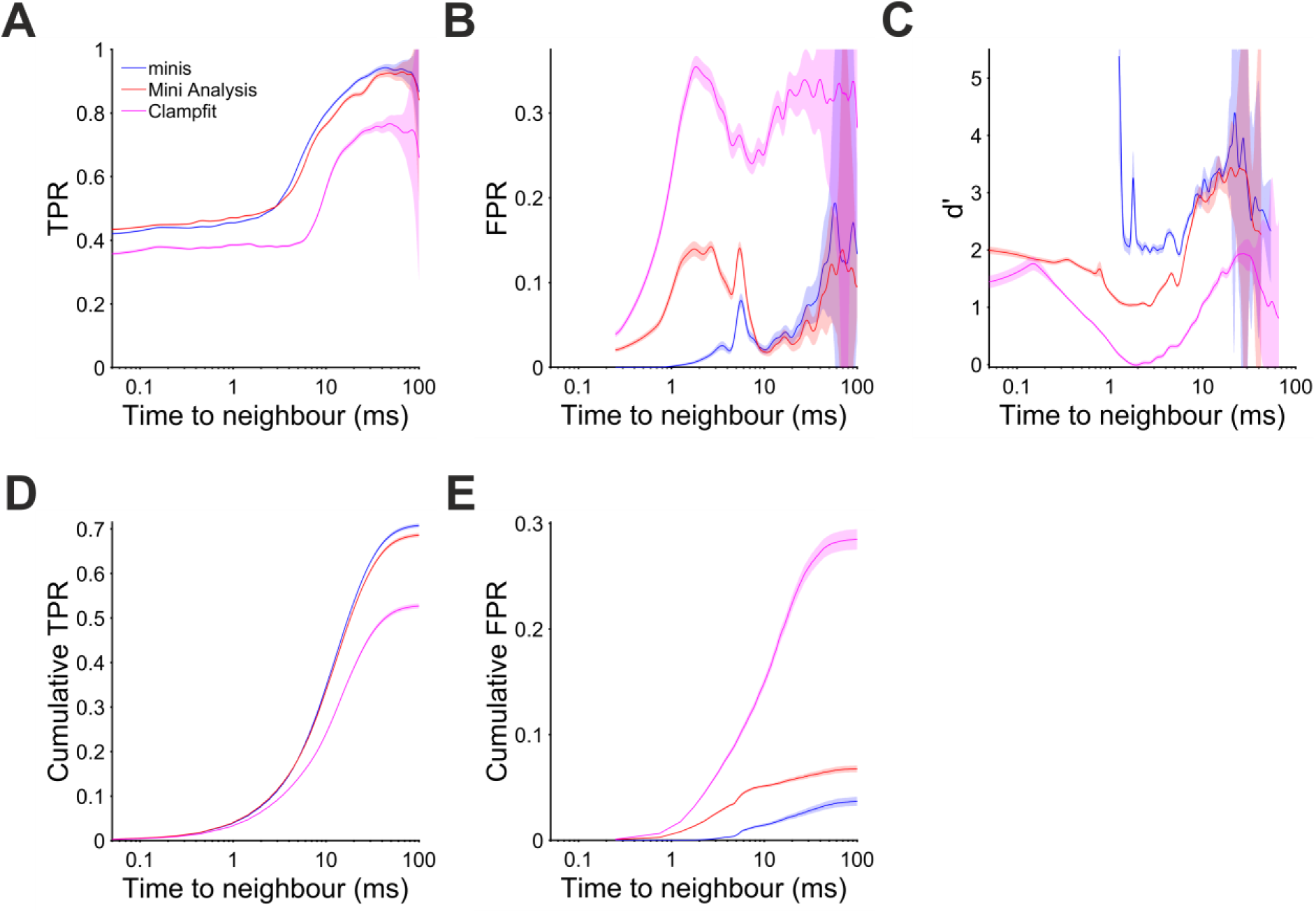
Performance when detecting moderately-sized (∼0.3 mV) smEPSPs as a function of time to the nearest neighbour with realistic mini rates (20, 40, 80 minis/s). Paler shaded colours indicate 95% confidence intervals. (A) True positive rate (TPR). (B) False positive rate (FPR). (C) Sensitivity index d’ (undefined but arbitrarily high for TPR > 0 and FPR = 0, e.g. for inter-mini intervals below 1 ms, for ‘minis’ algorithm). (D) Cumulative TPR. (E) Cumulative FPR.

**Supplementary Figure 2:**
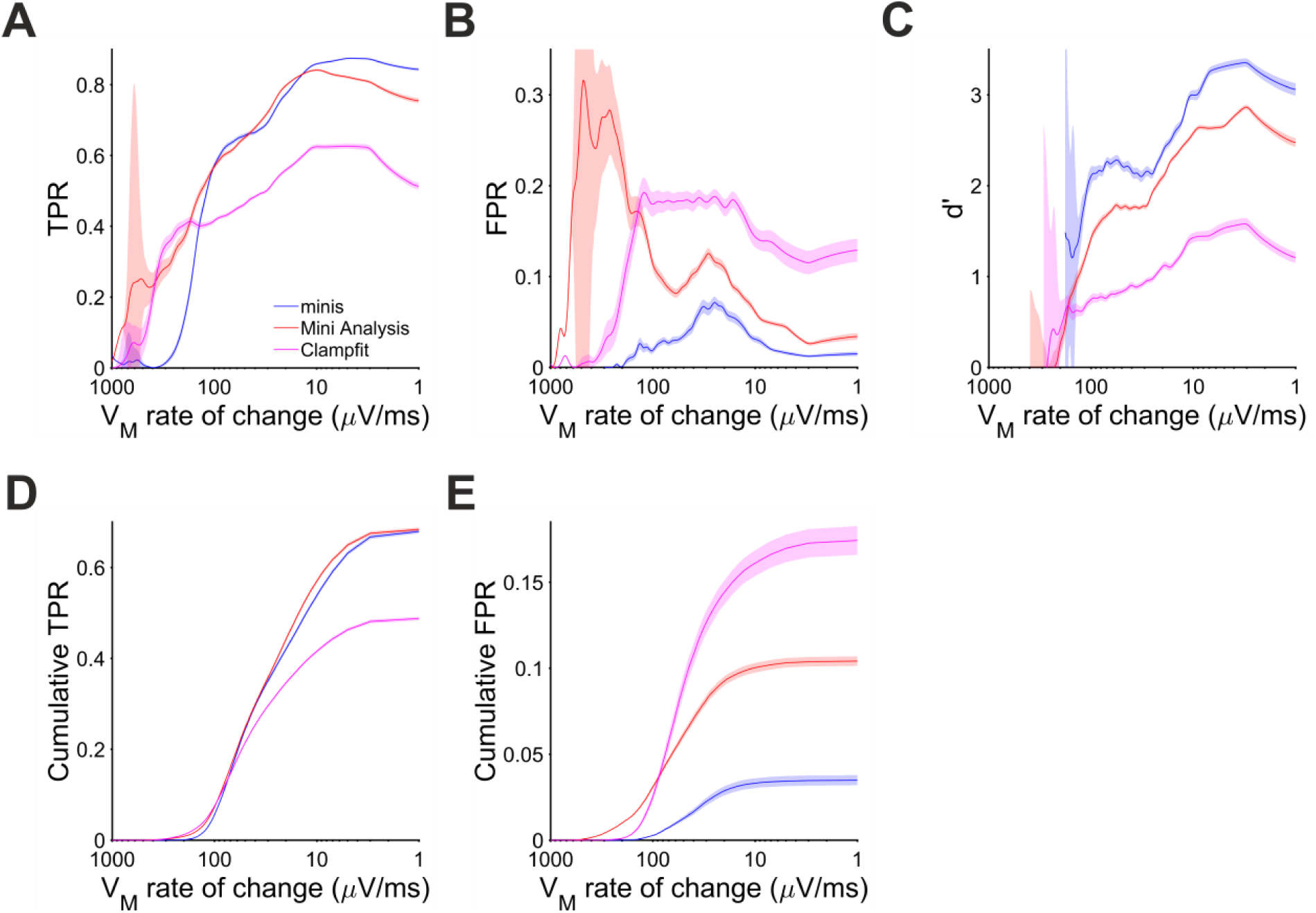
Performance when detecting moderately-sized (∼0.3 mV) smEPSPs on the rising trend V_m_ phases with realistic minis’ rates (20, 40, 80 minis/s, as in Fig. 9), as a function of the rate of change of V_m_. Pale shaded colours indicate 95% confidence intervals. (A) True positive rate (TPR) (B) False positive rate (FPR). (C) Sensitivity index d’. (D) Cumulative TPR. (E) Cumulative FPR.

**Supplementary Figure 3:**
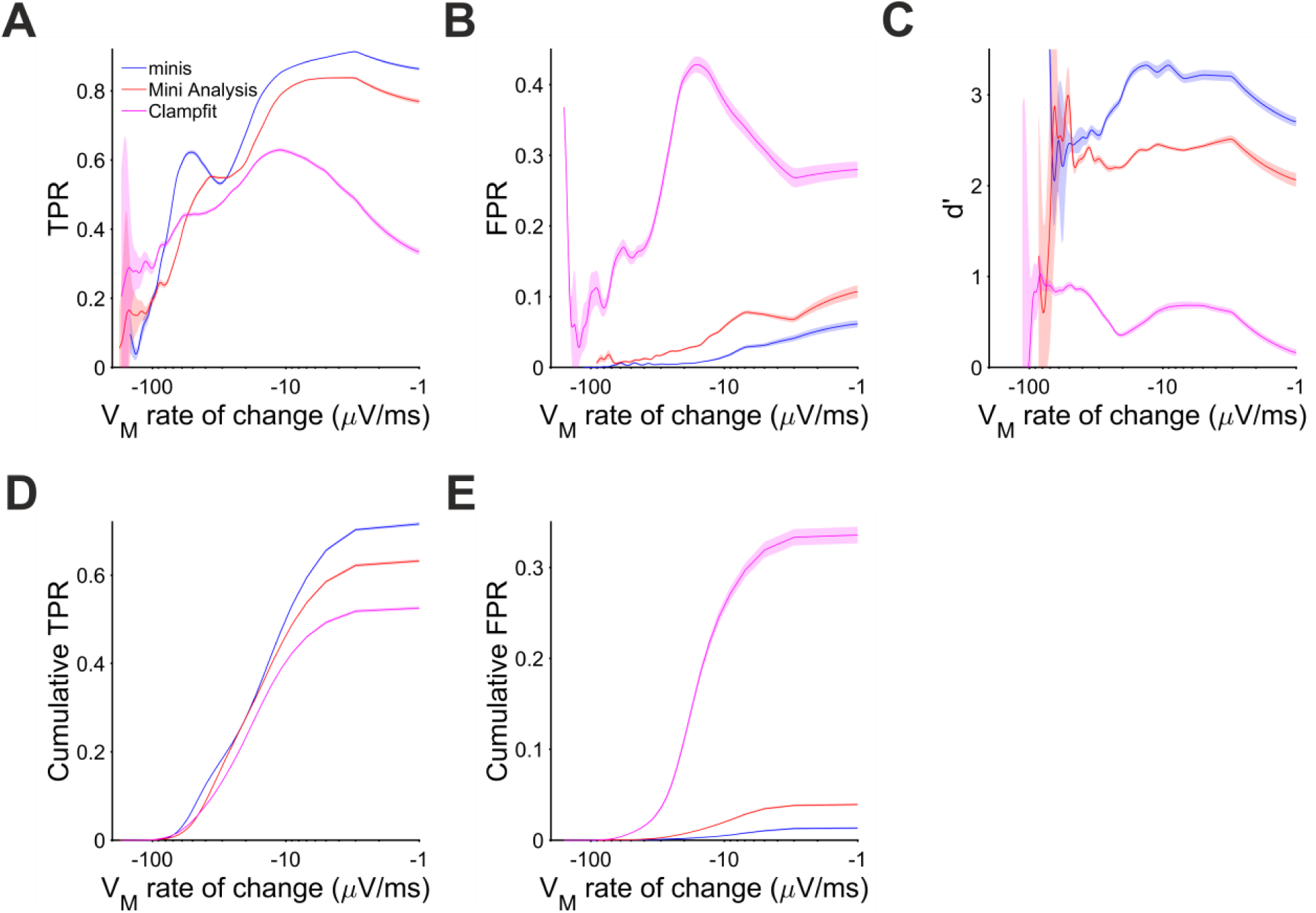
Performance when detecting moderately-sized (∼0.3 mV) smEPSPs on the membrane potential decay phase with realistic minis rates (20, 40, 80 minis/s, as in Figs. 9 and 10). Pale shaded colours indicate 95% confidence intervals. (A) True positive rate (TPR) when detecting moderately-sized smEPSPs on the V_m_ decay phase as a function of the V_m_ rate of change. (B) False positive rate (FPR). (C) Sensitivity index d’. (D) Cumulative TPR. (E) Cumulative FPR.

**Supplementary Table 1:**
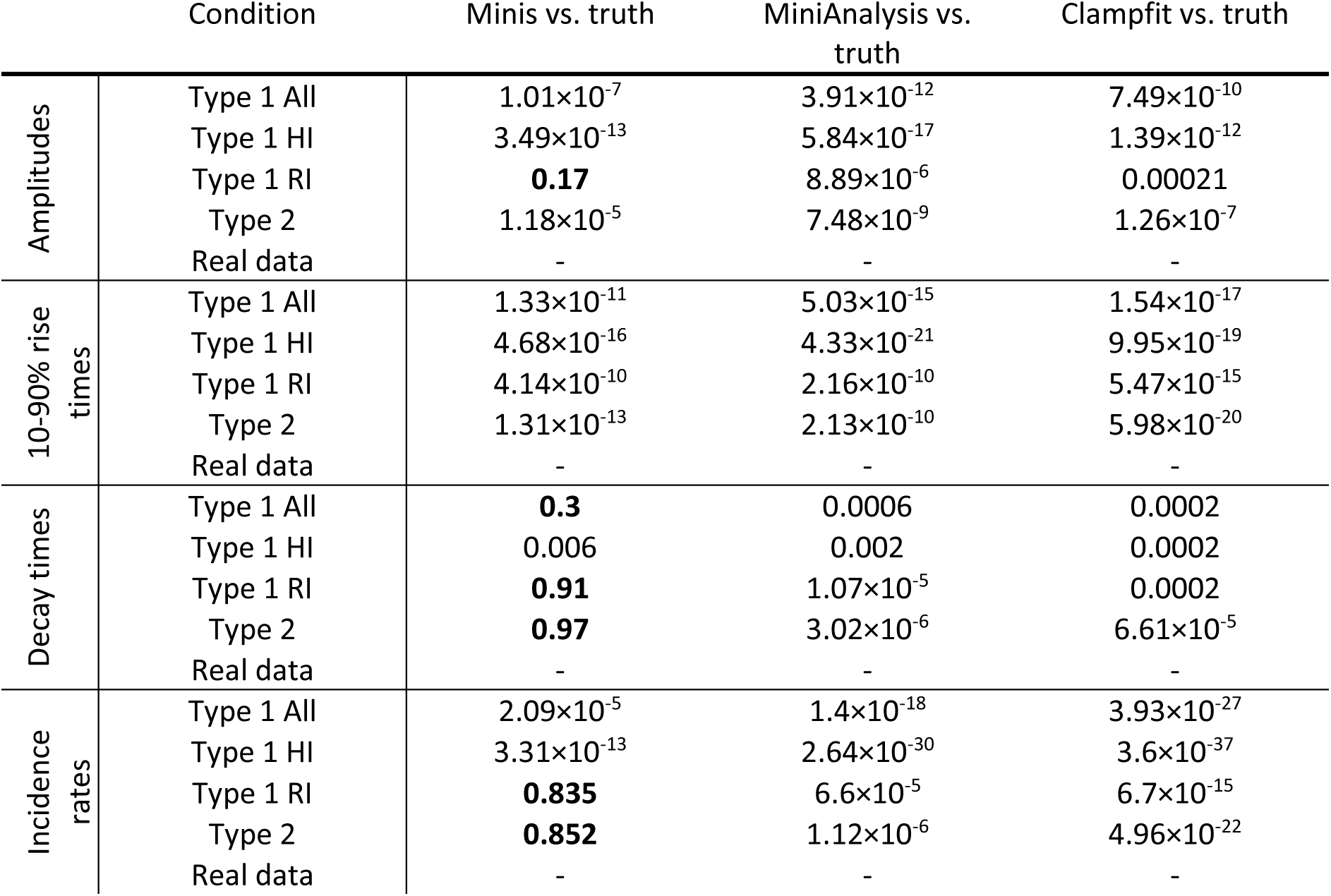
T-test p-values for mean amplitude, 10-90% rise time, decay time, and incidence rate values of detected minis using different algorithms. Repeated measures t-tests were used when comparing performance of two algorithms and single sample t-tests were used when comparing mean values with ground truth (subtracting ground truth values). Abbreviations are the same as in Table 1.

## References

Agmon-Snir H, Segev I. 1993. Signal delay and input synchronization in passive dendritic structures. J Neurophysiol 70:2066–2085. doi:10.1152/jn.1993.70.5.2066

Brown TH, Wong RKS, Prince DA. 1979. Spontaneous miniature synaptic potentials in hippocampal neurons. Brain Res 177:194–199. 10.1016/0006-8993(79)90931-4

Clements JD, Bekkers JM. 1997. Detection of spontaneous synaptic events with an optimally scaled template. Biophys J 73:220–229. 10.1016/S0006-3495(97)78062-7

De Koninck Y, Mody I. 1994. Noise analysis of miniature IPSCs in adult rat brain slices: properties and modulation of synaptic GABAA receptor channels. J Neurophysiol 71:1318–1335. doi:10.1152/jn.1994.71.4.1318

del Castillo J, Katz B. 1954. Quantal components of the end-plate potential. J Physiol 124:560–573. 10.1113/jphysiol.1954.sp005129

Dervinis M. 2024a. https://gin.g-node.org/dervinism/minis-benchmarking-data2. GIN. doi:10.12751/g-node.mvsy4j

Dervinis M. 2024b. https://github.com/dervinism/minis-benchmarking. GitHub. doi:10.5281/zenodo.14278784

Dervinis M. 2024c. https://github.com/dervinism/minis. GitHub. doi:10.5281/zenodo.14573132

Dervinis M, Major G. 2024. Novel quantal analysis method reveals conservation of average excitatory synaptic charge across cortical pyramidal neurons of different sizes. bioRxiv. doi:10.1101/2024.07.05.602190

El Khoueiry C, Alba-Delgado C, Antri M, Gutierrez-Mecinas M, Todd AJ, Artola A, Dallel R. 2022. GABAA and Glycine Receptor-Mediated Inhibitory Synaptic Transmission onto Adult Rat Lamina IIi PKCγ-Interneurons: Pharmacological but Not Anatomical Specialization. Cells 11. doi:10.3390/cells11081356

Eyal G, Verhoog MB, Testa-Silva G, Deitcher Y, Benavides-Piccione R, DeFelipe J, de Kock CPJ, Mansvelder HD, Segev I. 2018. Human Cortical Pyramidal Neurons: From Spines to Spikes via Models. Front Cell Neurosci.

Fatt P, Katz B. 1952. Spontaneous subthreshold activity at motor nerve endings. J Physiol 117:109–128.

Glasgow SD, McPhedrain R, Madranges JF, Kennedy TE, Ruthazer ES. 2019. Approaches and Limitations in the Investigation of Synaptic Transmission and Plasticity. Front Synaptic Neurosci 11. doi:10.3389/fnsyn.2019.00020

Holler S, Köstinger G, Martin KAC, Schuhknecht GFP, Stratford KJ. 2021. Structure and function of a neocortical synapse. Nature 591:111–116. doi:10.1038/s41586-020-03134-2

Hwang TN, Copenhagen DR. 1999. Automatic detection, characterization, and discrimination of kinetically distinct spontaneous synaptic events. J Neurosci Methods 92:65–73. 10.1016/S0165-0270(99)00095-3

Isaac JTR, Nicoll RA, Malenka RC. 1995. Evidence for silent synapses: Implications for the expression of LTP. Neuron 15:427–434. 10.1016/0896-6273(95)90046-2

Isaacson JS, Walmsley B. 1995. Counting quanta: Direct measurements of transmitter release at a central synapse. Neuron 15:875–884. doi:10.1016/0896-6273(95)90178-7

Larkman AU. 1991. Dendritic morphology of pyramidal neurones of the visual cortex of the rat: III. Spine distributions. Journal of Comparative Neurology 306:332–343. 10.1002/cne.903060209

Larkum ME, Nevian T, Sandler M, Polsky A, Schiller J. 2009. Synaptic Integration in Tuft Dendrites of Layer 5 Pyramidal Neurons: A New Unifying Principle. Science (1979) **325**:756–760. doi:10.1126/science.1171958

Liao D, Hessler NA, Malinow R. 1995. Activation of postsynaptically silent synapses during pairing-induced LTP in CA1 region of hippocampal slice. Nature 375:400–404. doi:10.1038/375400a0

Liao D, Jones A, Malinow R. 1992. Direct measurement of quantal changes underlying long-term potentiation in CA1 hippocampus. Neuron 9:1089–1097. 10.1016/0896-6273(92)90068-O

Macmillan NA, Creelman CD. 2005. Detection theory: A user’s guide, 2nd ed., Detection theory: A user’s guide, 2nd ed. Mahwah, NJ, US: Lawrence Erlbaum Associates Publishers.

Major G, Evans JD, Jack JJ. 1993. Solutions for transients in arbitrarily branching cables: I. Voltage recording with a somatic shunt. Biophys J 65:423–449. doi:10.1016/S0006-3495(93)81037-3

Major G, Larkman AU, Jonas P, Sakmann B, Jack JJ. 1994. Detailed passive cable models of whole-cell recorded CA3 pyramidal neurons in rat hippocampal slices. The Journal of Neuroscience 14:4613. doi:10.1523/JNEUROSCI.14-08-04613.1994

Major G, Larkum ME, Schiller J. 2013. Active Properties of Neocortical Pyramidal Neuron Dendrites. Annu Rev Neurosci 36:1–24. doi:10.1146/annurev-neuro-062111-150343

Motta A, Berning M, Boergens KM, Staffler B, Beining M, Loomba S, Hennig P, Wissler H, Helmstaedter M. 2019. Dense connectomic reconstruction in layer 4 of the somatosensory cortex. Science (1979) 366:eaay3134. doi:10.1126/science.aay3134

Murthy VN, Stevens CF. 1999. Reversal of synaptic vesicle docking at central synapses. Nat Neurosci 2:503–507. doi:10.1038/9149

Nevian T, Larkum ME, Polsky A, Schiller J. 2007. Properties of basal dendrites of layer 5 pyramidal neurons: a direct patch-clamp recording study. Nat Neurosci 10:206–214. doi:10.1038/nn1826

Rall W. 1977. Core Conductor Theory and Cable Properties of Neurons In: Kandel ER, Brookhardt JM, Mountcastle VM, editors. Handbook of Physiology, Cellular Biology of Neurons. Bethesda, MD: American Physiological Society. pp. 39–97.

Rall W. 1967. Distinguishing theoretical synaptic potentials computed for different soma-dendritic distributions of synaptic input. J Neurophysiol 30:1138–1168. doi:10.1152/jn.1967.30.5.1138

Rinzel J, Rall W. 1974. Transient Response in a Dendritic Neuron Model for Current Injected at One Branch. Biophys J 14:759–790. 10.1016/S0006-3495(74)85948-5

Sara Y, Bal M, Adachi M, Monteggia LM, Kavalali ET. 2011. Use-Dependent AMPA Receptor Block Reveals Segregation of Spontaneous and Evoked Glutamatergic Neurotransmission. The Journal of Neuroscience 31:5378. doi:10.1523/JNEUROSCI.5234-10.2011

Segal M. 2010. Dendritic spines, synaptic plasticity and neuronal survival: activity shapes dendritic spines to enhance neuronal viability. European Journal of Neuroscience 31:2178–2184. doi: 10.1111/j.1460-9568.2010.07270.x

Shi Y, Nenadic Z, Xu X. 2010. Novel Use of Matched Filtering for Synaptic Event Detection and Extraction. PLoS One 5:e15517-.

Stuart G, Spruston N. 1998. Determinants of Voltage Attenuation in Neocortical Pyramidal Neuron Dendrites. The Journal of Neuroscience 18:3501–3510. doi:10.1523/JNEUROSCI.18-10-03501.1998

Wall MJ, Usowicz MM. 1998. Development of the quantal properties of evoked and spontaneous synaptic currents at a brain synapse. Nat Neurosci 1:675–682. doi:10.1038/3677

Williams SR, Mitchell SJ. 2008. Direct measurement of somatic voltage clamp errors in central neurons. Nat Neurosci 11:790–798. doi:10.1038/nn.2137

Williams SR, Stuart GJ. 2002. Dependence of EPSP Efficacy on Synapse Location in Neocortical Pyramidal Neurons. Science (1979) **295**:1907–1910. doi:10.1126/science.1067903

## Supplementary References

del Castillo J, Katz B. 1954. Quantal components of the end-plate potential. J Physiol 124:560–573. doi: 10.1113/jphysiol.1954.sp005129

Dervinis M. 2024c. https://github.com/dervinism/minis. GitHub. doi:10.5281/zenodo.14573132

Larkum ME, Nevian T, Sandler M, Polsky A, Schiller J. 2009. Synaptic Integration in Tuft Dendrites of Layer 5 Pyramidal Neurons: A New Unifying Principle. Science (1979) 325:756–760. doi:10.1126/science.1171958

Segal M. 2010. Dendritic spines, synaptic plasticity and neuronal survival: activity shapes dendritic spines to enhance neuronal viability. European Journal of Neuroscience 31:2178–2184. 10.1111/j.1460-9568.2010.07270.x

